# A Multidisciplinary Approach to Explain Biological Aging and Longevity

**DOI:** 10.1101/045633

**Authors:** Brett N. Augsburger

**Affiliations:** Department of Pathobiology Auburn University

## Abstract

Scientists have been unable to reach a consensus on why organisms age and why they live as long as they do. Here, a multidisciplinary approach was taken in an attempt to understand the root causes of aging. Nonequilibrium thermodynamics may play a previously unappreciated role in determining longevity by governing the dynamics of degradation and renewal within biomolecular ensembles and dictating the inevitability of fidelity loss. The proposed model offers explanations for species longevity trends that have been previously unexplained and for aging-related observations that are considered paradoxical within current paradigms—for example, the elevated damage levels found even in youth within many long-lived species, such as the naked mole-rat. This framework questions whether declining selective pressure is the primary driver of aging, and challenges major tenets of the disposable soma theory. Unifying pertinent principles from diverse disciplines leads to a theoretical framework of biological aging with fewer anomalies, and may be useful in predicting outcomes of experimental attempts to modulate the aging phenotype.

## 1 Introduction

Many theories to explain why organisms age have been proposed. Concepts from evolutionary theory, genetics, biochemistry, and cellular and molecular biology are most often used as the basis of these theories. Despite the fact that these efforts have resulted in a multitude of theories, each with serious anomalies, the focus over the last half century has remained in these areas—more fundamental physical law has been under or incorrectly applied, or simply ignored altogether. Notwithstanding the fact that deterioration is implicated nearly universally in the aging process, the connection to nonequilibrium thermodynamics and entropy production has not been firmly established and is infrequently mentioned. There have been a few notable exceptions; for example, Hayflick contends that entropy alone is sufficient to explain biological aging (Hayflick 2000; Hayflick 2004; Hayflick 2007b; Hayflick 2007a).

The second law of thermodynamics (hereafter abbreviated to “second law”) stipulates that all energy, regardless of form, has a propensity to transition from a localized state to one that is more spread out—i.e. dispersed in space—at a rate determined by the ability of contributing external factors to counteract this tendency. Entropy is a measure of this "spreading out" of energy. In any system not in thermodynamic equilibrium, the second law stipulates that entropy will be produced by means of irreversible processes. This entropy is referred to as "internal entropy". A nonequilibrium system will continue to produce internal entropy indefinitely until equilibrium is reached, resulting in a transition from a higher concentration of molecular bond energy to a lower bond energy concentration (Demirel 2014).

It has been argued that the second law only relates to isolated systems and that since organisms are open systems the second law does not apply (Mitteldorf 2010). This is false—the second law is universally applicable and always satisfied (Kondepudi and Prigogine 2014). According to modern nonequilibrium thermodynamics, the second law describes the tendency for internal entropy to be produced by any system that is not in equilibrium. Clearly, organisms are not in thermodynamic equilibrium and therefore all living organisms continuously produce internal entropy. Despite existing in a nonequilibrium state, individual organisms resist the decay towards equilibrium, which eventually results in death, long enough to allow them to mature and reproduce. The term “longevity”, as used here, refers to the average length of time that individuals of a group live under ideal conditions (which can range between hours to centuries, depending on the species).

Organisms combat entropy increases resulting from internal entropy production by exchanging heat and other forms of energy with their surroundings and importing/exporting negative or positive entropy in the form of metabolites and catabolites. Some scientists have suggested that, using these means, any entropy increase in an organism can always be counteracted without permanent negative repercussions to the individual; from this, some have concluded that there is no thermodynamic stipulation for aging to occur and no role for thermodynamics in explaining aging. This is a rather obvious *non sequitur*—yet this notion has been perpetuated in the aging literature, both explicitly and implicitly (Kirkwood 1999; Mitteldorf 2010; Trindade et al. 2013). Here it is demonstrated why this inference is a logical fallacy and how it neglects to consider critical effects of thermodynamic phenomena within an organism— particularly the influence of internal entropy production on the flow of biomolecular-encoded information over time and the dynamics of degradation and renewal within biomolecular ensembles.

## 2 Nonequilibrium Thermodynamics Stipulates Biomolecular Damage

Thermodynamic equilibrium is characterized by the total absence of thermodynamic potentials within a system and the minimization of free energy. As the atomic arrangement of biomolecules within an organism fails to maximize free energy^1^—energy remains highly concentrated due to the chemical bonds holding biomolecules together—the second law stipulates that irreversible processes driving the system in the direction of equilibrium must occur and impose insults on biomolecular structure. The resulting thermodynamic fluxes (thermal, chemical, diffusive, mechanical, electrical, etc.) within an organism contain significant spatial and temporal heterogeneity at both the mesoscopic and macroscopic levels, providing a multitude of opportunities for biomolecular interactions leading to undesirable structural states. Examples include: mechanical force-based unfolding of proteins; unzipping or shearing of nucleic acids; protein folding alterations and undesirable protein associations due to crowding (Zhou 2010); DNA hydrolysis, oxidation, and methylation reactions (Lindahl 1993); denaturation of DNA and protein from excessive temperature; and disruption of hydrophobic interactions leading to altered protein structure. It should be apparent that while all else being equal it is beneficial for an organism to minimize the probabilities of these occurrences, it is impossible to reduce them to zero. Organisms counteract this degradation by producing negative entropy. In terms of preservation of overall biomolecular^2^ integrity, an organism could be viewed as a near steady state nonequilibrium system—at least if considered over a snapshot-in-time that is short compared to total lifespan.

### 2.1 A Model System for Analyzing Thermodynamically Derived Biomolecular Degradation

A model system is defined below to demonstrate the degradative effects of thermodynamic chemical forces on biomolecules. The analysis focuses on a single type of biomolecule existing in a fixed volume of cytosol. The biomolecule of interest could be any expressed protein, synthesized lipid or other biomolecule produced by the organism's cellular machinery. For this analysis, the temperature and pressure of the system are in equilibrium with the surroundings and equivalent to physiological values. The concentrations of all molecules apart from the biomolecule of interest are held constant by chemiostats. The system remains in thermomechanical equilibrium at all times. Chemical reactions not involving the biomolecule of interest are inhibited (reaction rates and affinities are zeroed), as are all biomolecular repair and replacement mechanisms. At time *t*_0_, the biomolecule of interest is in a state where all of its constituents are maximally capable of performing its intrinsic biological function.

In accordance with the second law, the described system will produce internal entropy and transition irreversibly from an initial state at time *t*_0_ through a very large number of nonequilibrium states until all chemical reaction affinities are reduced to zero. Since the system as defined has no means of counteracting internal entropy production, the second law requires that the only stable or steady state is the chemical equilibrium state. Although some reaction affinities may be low, even the most improbably reactions must have nonzero reaction rates. During the progression towards chemical equilibrium, the biomolecule can exist in a very large number of alternative internal states representing various degradative arrangements. The transitions between internal states can be characterized by reactions of the form

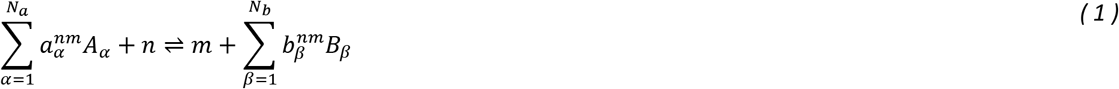

where *n* and *m* are the initial and new protein internal states, 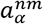 and 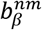 are the number of molecules of reactant *(A_α_)* or product (*B_β_*) involved in the reaction, and *N_a_* and *N_b_* are the number of different reactants and products involved. Until the system reaches equilibrium, these reactions will occur and produce internal entropy. The rate of internal entropy production *d_i_S/dt* at any time *t* up until equilibrium can be expressed as

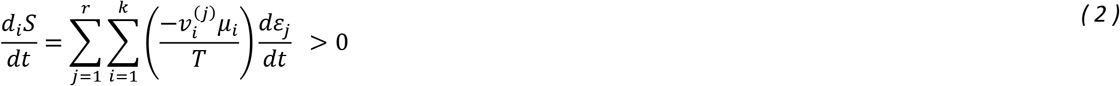

where *k* is the number of chemical species involved in a particular reaction, *r* is the total number of reactions taking place in the system, 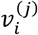 are the stoichiometric coefficients, *µ_i_* are the chemical potentials, and *dε_j_/dt* represents the reaction velocity at time *t*.

At thermodynamic equilibrium, both the reaction velocity and the reaction affinity 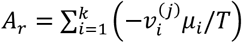 will be zero. This represents a state where all of the examined biomolecule have fully degraded and internal entropy production has ceased.

The total entropy increase in the system at any time *t* is

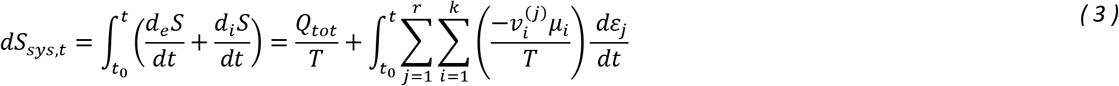

Here, *d_e_S/dt* represents the rate of entropy gain/loss in the system due to the exchange of energy with the surroundings (heat flowing into or out of the system). *Q_tot_* represents the total heat that has been transferred to/from the system between time *t*_0_ and time *t*. Utilizing the change in system entropy *dS_sys,t_* for any time *t* and the increase in system entropy corresponding to the equilibrium condition *dS_sys,max_*, we define a new parameter termed “degradation state” *D* to represent the degree to which the biomolecule under examination has degraded

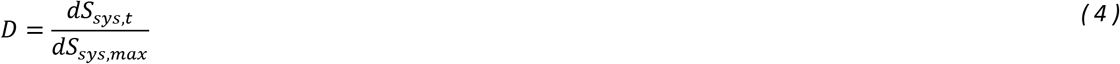

A *D* of 1 corresponds to full degradation, while a value of 0 is representative of a pool of fully intact biomolecules.

Of course, degradative internal entropy production within a living organism is not limited to chemical reactions but also includes contributions from heat, mass, and momentum transfer as well as electrical, magnetic, diffusion and other effects. Modern nonequilibrium thermodynamic theory (and the second law in particular) can be utilized to similarly model each of these factors, establishing an arrow of time stipulating that the future is distinguishable from the past by an ever-increasing quantity of total internal entropy produced.

### 2.2 Preservation of Steady State Nonequilibrium within a Biomolecular System

For homeostasis—i.e. steady state—to be preserved and to avoid transitioning to an equilibrium state with maximum disorder, degradation must be combatted by biological mechanisms capable of producing sufficient negative entropy to fully offset the internal entropy produced, such that the total system entropy is unchanged:

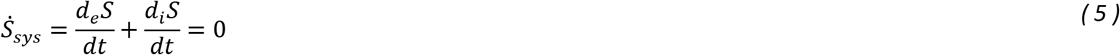

Since the second law stipulates that, 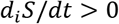 in order to maintain a steady state

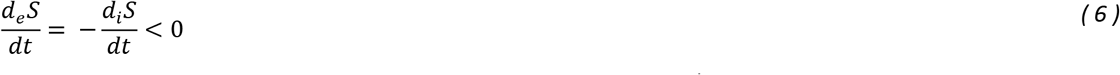

Suppose that a renewal mechanism is incorporated which exchanges *Ṅ_rc_* moles s^-1^ of degraded biomolecules with newly expressed or fully repaired biomolecules. A steady state is maintained if

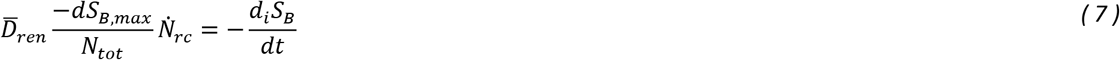

where 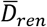 represents the average degradation state of a renewed biomolecule and *N_tot_* is the total quantity of the biomolecule of interest in moles. (The rate of internal entropy production will be denoted by *Ṡ_i_* instead of *d_i_S_B_/dt* hereafter for purposes of clarity. Also, note that *S_B_* and *S_B,max_* refer to the fraction of the entropy associated with the total population of the biomolecule being analyzed.) Eq. *(7)* can be rearranged to solve for renewal rate during steady state.

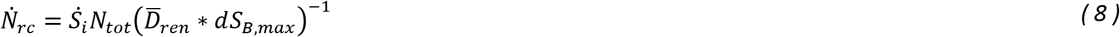

The degradation state *D* of a biomolecular ensemble specifies the level of degradation of the average biomolecule but it does not indicate how well biomolecules perform at that degradation state. The term “biomolecular performance” *P* will be used to quantify the relative ability of a biomolecular ensemble to perform its intrinsic biological function(s). A value for *P* of 1 indicates that the average biomolecule in an ensemble is able to perform 100% of its intrinsic biological function(s), or in other words, the ensemble will perform as if all biomolecules were in ideal condition. A *P* of 0 denotes that the average biomolecule is degraded to the extent that it is unable to perform any of its intrinsic biological function.

### 2.3 Relating Biomolecular Performance to Degradation State

Biomolecular insults are inevitable occurrences; for this reason, biomolecules might be expected to retain the ability to perform their intrinsic biological function even when some level of damage is present. If a biomolecule did not have this capability, only a very small percentage of biomolecules within a pool would likely be functional at any given time. It is hypothesized here that the falloff in *P* with increasing *D* could be approximated for many biomolecules by a logistic function. The following 4-parameter logistic function describing the relationship between *P* and *D* produced an acceptable fit to the empirical data examined here:

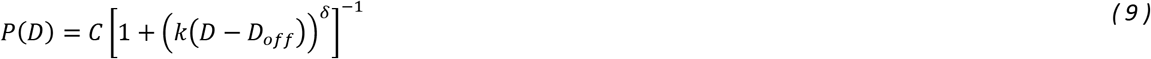

where *C, k, D_off_*, and *δ* are the fitting parameters.

### 2.4 Biomolecular Work Rate is a Primary Determinant of the Rate of Degradative Internal Entropy Production

A transfer of energy occurs when a biomolecule performs its biological function, resulting in a period of time when the concentration of energy in close proximity to the biomolecule is significantly elevated compared to the resting (or nonworking) state. This generates higher thermodynamic potentials in the participating biomolecules and therefore elevates *Ṡ_i_*— i.e. the rate of damage increases transiently during biological processes. This work-related damage is one and the same as that attributed to the inherent imperfectness of all biological processes, as discussed by Gladyshev (2013; 2014; 2016). Since it is impossible for a biological process to occur without a transfer of energy and hence increased thermodynamic potentials, no biological process can be perfect and stochastic deleterious biomolecular insults will inevitably occur at a higher rate while biological processes are in progress. All else being equal, a biomolecular pool that is inactive (not performing any intrinsic biological functions) will exhibit lower 5¿ than one in which biomolecules spend more time in the active (working) state.

### 2.5 Relating Internal Entropy Production Rate to Degradation State

The fraction of time that a biomolecule performs doing work compared to when the biomolecule is inactive is an important factor in determining the rate of internal entropy production. In addition, biomolecules existing at a lower degradation state, and thus exhibiting higher performance, are capable of performing more work within a given period of time and in doing so will transfer energy at a higher rate during a biological process, thereby leading to higher rates of damaging internal entropy production.

It is hypothesized here that, for a given biomolecule, the amount of internal entropy produced per unit of work produced is similar across a relatively large range of physiologically useful *P/D* values yet not without variance. The following equation summarizes this model of the rate of internal entropy production within a biomolecular ensemble resulting from work

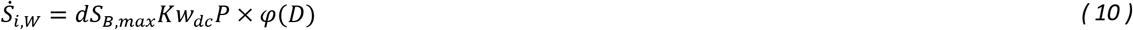

*w_dc_* is the work duty cycle (the relative amount of time biomolecules spend doing work, a value between 0 and 1) and *K* is a scaling factor. The function *φ(D)* compensates for any non-linearities in internal entropy production rate across degradation state.

Internal entropy is also produced in the static (non-working) conditions. In the described model system, the magnitude of the thermodynamic forces on the biomolecules of interest during non-working conditions should remain essentially constant and be independent of *P* or *D*. The rate of internal entropy production during non-working conditions is represented here by *L* × φ^(*D*)^, where *L* is a scaling constant for non-working conditions. The total internal entropy production is thus

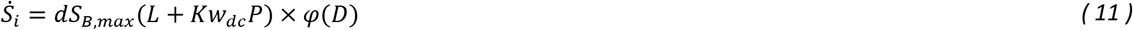

From Eq. *(4)* we know that *Ṡ_i_ = dS_B,max_Ḋ*. Combining this with the previous equation leads to an expression for the rate of change of the degradation state

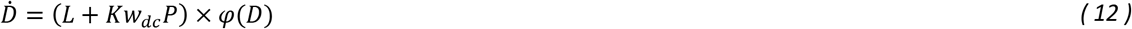

For a system consisting of molecules with various degrees of deterioration, *Ḋ* can be determined by considering the number of molecules existing at each represented equivalent^3^ *D* and the corresponding equivalent *Ḋ(Ḋ_D_*, calculated using Eq. *(12)*)

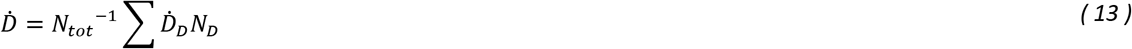

### 2.6 Incorporating Biomolecular Repair and Replacement Mechanisms

Two different renewal scenarios will be considered. For both scenarios, it is assumed that the renewal process returns the biomolecules to their ideal functional condition (equivalent *D* of 0). In the first and simplest scenario, biomolecules are chosen for renewal at random; i.e. biomolecules in perfect condition (high equivalent *D)* have the same probability of being renewed as those in poor condition (low equivalent *D*). In this case, 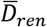 is equal to *D*.

To reduce the resources directed towards renewal and to maximize the return on investment into renewal, the renewal machinery should be able to distinguish more degraded biomolecules from those that are less degraded. In a second renewal scenario, this ability is introduced. Here, the molecules chosen for renewal are those whose equivalent *P* places them in the lowest *“m”* fraction of all molecules within the ensemble.

Ideally, any molecular damage that decreases *P* would be detectable by cellular machinery. In practice, this is not possible to achieve with perfection. The only way to definitively confirm that a biomolecule is in ideal condition is to observe the biomolecule while it is performing work, and verify that the magnitude and quality of the work performed are consistent with a biomolecule in ideal condition; this requires faultless labelling and work-measurement systems which in turn would require their own such systems, etc. Perhaps a more practical way of identifying biomolecular damage is by the direct recognition of molecular alterations with generic signatures. For example, oxidative damage can readily be recognized with universal mechanisms. However, this is only one type of damage and these mechanisms are inherently imperfect. As the number of possible molecular alterations that could affect biomolecular equivalent *P* is essentially unlimited, recognizing every one of these alterations—and doing so with perfect precision — is simply not possible.

### 2.7 Balanced Biomolecular Renewal and Degradation: Numerical Approximation of the Steady State Solution

A biomolecular ensemble undergoing continuous renewal and degradation will consist of biomolecules distributed across a range of equivalent *P* / *D* values. As it is cumbersome to find a closed-form solution of this model, here an approximate solution is calculated numerically using iteration. By combining Eqs. *(4), (8)* and *(13)*, it can be shown that a steady state will be maintained within an biomolecular ensemble (i.e. *D* will remain constant) when the following condition is satisfied

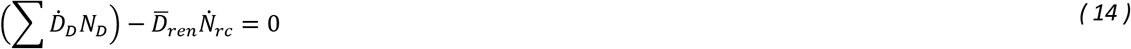

That is to say that any positive contribution to *D* resulting from internal entropy production must be counterbalanced with an equivalent negative contribution from renewal. The magnitude of the left-hand side of Eq. *(14)* is therefore a measure of solution convergence.

In the numerical analysis undertaken here, a fixed number of biomolecules are distributed across a range of *D* values. The initial distribution at *t* = 0 is not important. For all values of *D* other than 0 and for each time point +Δt, the number of molecules at each equivalent *D* value represented, *N_D_*, is calculated

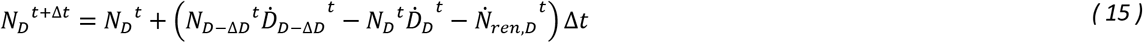

where *Ṅ_ren,D_* is the rate at which molecules of a particular equivalent *D* are renewed. This value is calculated according to the renewal scenario (from section 2.6) that is specified and using the distribution of biomolecules across equivalent degradation states at time *t. Ḋ_D_* and *Ḋ_D-ΔD_* are calculated per Eq. *(12)* for each represented *D*.

For the present analyses, it is assumed that *φ*(*D*) = 1. This is done because (1) determining *φ*D** would require knowing the system entropy for different *P* values and this is a difficult property to measure empirically, (2) it is reasonable to hypothesize that *φ* is relatively constant within the utilized functional ranges found within organisms since any significant perturbations in *φ*D** would indicate either an undesirable vulnerability or unnecessary stability for the sake of some other factor, and (3) the performance/renewal rate insights that follow remain valid because *D* and *Ṡ* are intermediate variables in these calculations (the effects of *φ* will effectively cancel out). *D* and *Ṡ_t_* have been retained in the derivations because conceptually they help to convey a more thorough understanding of the underlying physical phenomenon at play.

For the case where *D* = 0, the following is used in lieu of Eq. *(15)* (Note that all renewed molecules are initialized to *D* = 0.)

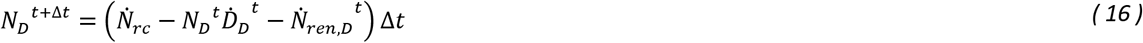

If the solution failed to converge to within a specified tolerance after a given number of iterations, the simulation was restarted using altered parameters to increase precision (e.g. larger vector of *D* values, reduced Δ*t*) at the expense of increased computational resources. These steps were repeated until convergence was reached.

### 2.8 Realizable Work Rate and Biological Power Density

The achievable rate of work per biomolecule *Ẇ_mact_* can be expressed as

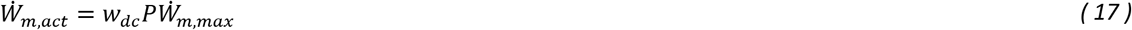

where *Ẇ_m,max_* is the maximum potential external work rate per molecule (corresponding to *P* = 1 and *w_dc_* = 1).

“Biological power density”, represented by *σ*, is defined as the volume-specific rate of external work (power per unit volume) realized in a biomolecular ensemble. In this context, “external work” refers to the sum of the biomechanical, biochemical and bioelectrical/electromagnetic work, representative of the intrinsic biological function of the biomolecule being analyzed, which is brought to bear on the immediate environment. Examples include the mechanical work produced by myosins, and the chemical and electrical work produced by cell surface receptors. Biological power density is given by

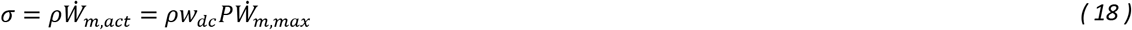

where *ρ* is the volumetric density (number of biomolecules per unit volume).

### 2.9 Demonstration of Model using Empirical Data

Given sufficient data describing the degradation characteristics of a biomolecule, the above model can be used to predict the following steady state properties: (a) the distribution of biomolecules across equivalent *P* values for any renewal rate; (b) how *P* varies with renewal rate; and (c) the renewal return on investment (ROI), calculated by dividing the work rate by renewal rate, as a function of *P*. The degradation dynamics of the biomolecule under work conditions must be known; i.e. how *P* diminishes over time while the biomolecules are actively performing work and no renewal is occurring. The degradation dynamics under non-working conditions are required if static degradation is to be considered.

Several factors make fluorescent biomolecules a good choice for demonstrating this model. For one, the energy they use to produce work comes from photons rather than a molecular energy source (e.g. ATP). Utilizing a readily available light source, a consistent and controllable energetic flux can be provided externally without disturbing the molecular composition of the system under analysis. This eliminates the need for chemiostats and the renewal/removal of molecular substrate. Since the work that is performed by fluorescent molecules is also realized in the form of photons, measuring the amount of work performed is straightforward.

*P* will decrease as fluorescent molecules absorb and emit photons. This is mostly a result of photobleaching, the irreversible loss of fluorescence due to photochemical modification of the fluorophore. Photobleaching is believed to be largely caused by interactions between excited fluorophores and triplet oxygen, leading to irreversible chemical damage to the fluorophore (Diaspro et al. 2006). The probability that a photon collision will result in excitation decreases with each subsequent photon absorbed. Since photobleaching damage is sustained almost exclusively when the fluorophore is in the excited state, incidental damage from photon collisions that fail to excite the fluorophore can be disregarded; i.e. provided that static degradation is slow compared to the rate of degradation from work, we can assume that the vast majority of any degradation observed is a direct result of producing work and not due to off target damage from the energy source.

All collisions between a photon and a fluorescent biomolecule will be considered work events, regardless of whether or not excitation occurs (i.e. even if no work is produced). The work event period is defined as the average amount of time for photon absorption and emission to occur. *w_dc_* can be precisely modulated by altering the intensity of the excitation source. The time to degrade under non-renewing conditions from an initial *P* of 1 to any given *P* can be calculated

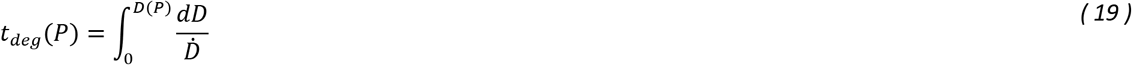

Using this equation together with previously published photobleaching data (Shaner et al. 2004), fitting parameters for Eq. *(9)* were determined for the fluorescent biomolecules eGFP, mRFP1, and mTangerine. Fig. 1A demonstrates the quality of the fit between the actual and predicted *P-t* curves for the parameter values calculated. Since the illumination intensity for each fluorophore (normalized such that each molecule emitted 1,000 photons/s at time *t* = 0) remained constant throughout each experiment, *w_dc_* can also be considered constant across each degradation curve (although not necessarily the same between fluorophores). The initial rate of work (photon emission) is considered to be representative of *P* = 1.

**Fig. 1.**
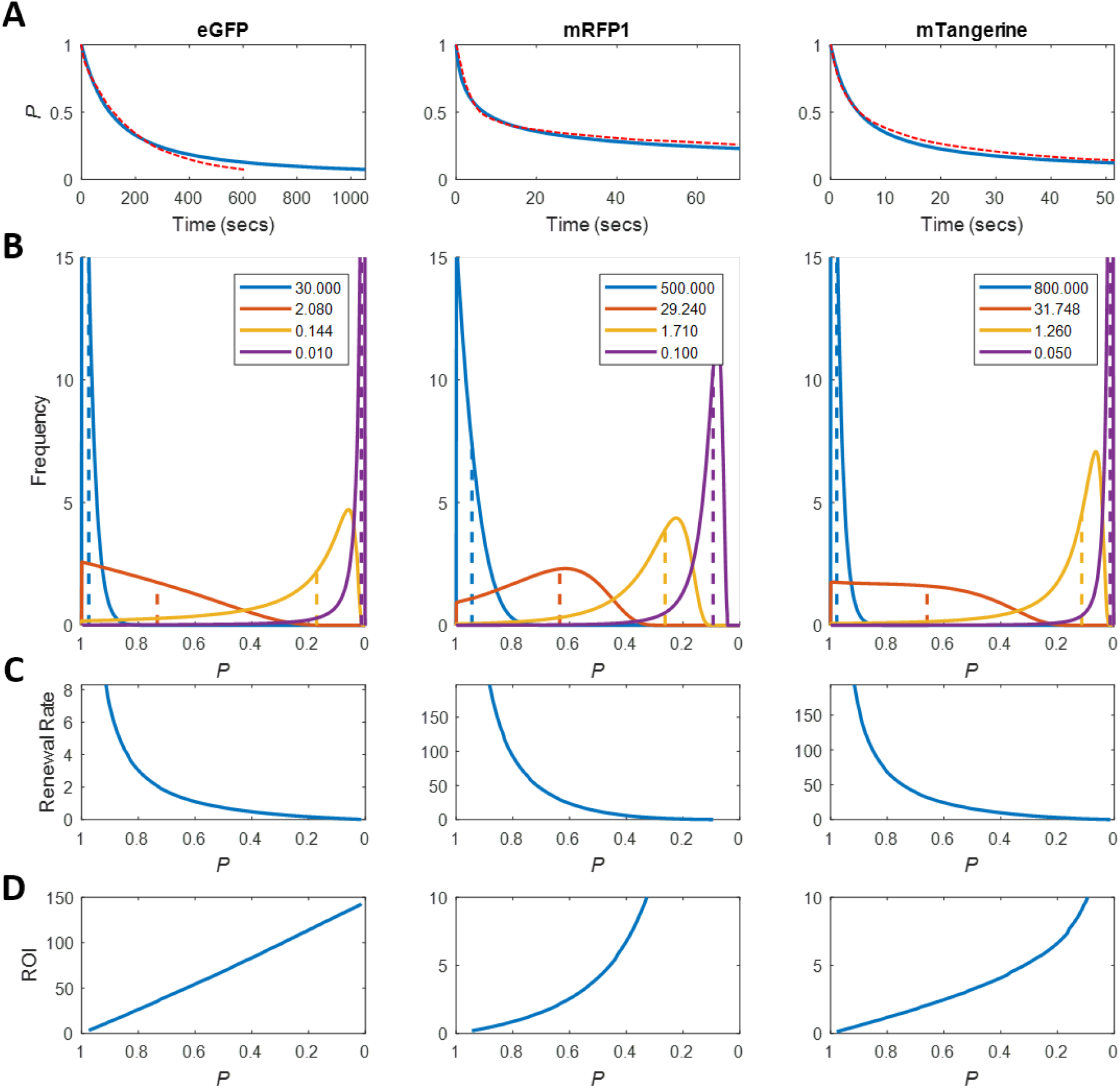
Relating degradation, renewal and performance characteristics in eGFP, mRFP1, and mTangerine populations. (a) Loss of P with time in a non-renewing population of eGFP, mRFP1, and mTangerine under work-producing conditions. Fitted curves (solid blue lines) versus empirical data (dashed red lines) from Shaner et al. (2004). All curves were fit to Eq. (9). Fitting parameters: eGFP — C = 2.7438, k = 0.1354, D_off_ = -7.3996, δ = 350.2; mRFP1 — C = 2.5414, k = 837.3, D_off_ = -0.0025, δ = 0.6; mTangerine — C = 1.3666, k = 38.2428, D_off_ = -0.0152, δ =1.85. (b) Simulation of distribution of biomolecules across equivalent P values for different rates of random renewal. Each simulation was iterated through until the solution converged to steady state. Renewal rates are in % of total population replaced per second. e.g. 500% indicates that the typical molecule is renewed 5 times every second. Dashed vertical lines indicate P of the ensemble (i.e. average equivalent P). (c) Relationship between P (ensemble) and renewal rate under random renewal conditions. D Renewal ROI versus P under random renewal. ROI is in % of W_m max_ / renewal rate. All simulations were performed with N_rc_ = 1e9.

### 2.10 High *P* is Energetically Costly. Perfect Fidelity in a Biomolecular Ensemble is Impossible.

In order to maintain a specific *P* in any biomolecular ensemble, renewal must take place; otherwise, *P* will decrease with time. Random renewal was simulated in eGFP, mRFP1, and mTangerine (using the fitting parameters found to approximate the deterioration characteristics of each fluorescent biomolecule). Only work-related degradation was considered. For each renewal rate, the simulation was allowed to iterate until convergence was achieved. It can be seen that the distribution of molecules across equivalent *P* values varies with renewal rate and is also dependent on the degradation characteristics of the biomolecule (Fig. 1B).

The rates of renewal required to maintain a particular *P* are shown in Fig. 1C. A common trend is a dramatic increase in renewal rate as *P* approaches 1, suggesting that achieving very high *P* requires an ever more costly investment in renewal. This trend is to be expected in any biomolecular ensemble and is due to several factors. Firstly, since the average equivalent *D* of a replaced molecule is lower when *P* is higher, the amount of negative entropy produced by each molecular renewal event will decrease with increasing *P*. Second, for a static *w_dc_*, the rate of internal entropy production increases with increasing *P* (as the amount of work produced increases) which further contributes to an increased renewal rate for larger *P* values. Furthermore, there is always a non-zero probability that some level of degradation will occur between the time a biomolecule is synthesized and the time it is renewed. Maintaining a *P* of 1 within any ensemble is therefore impossible.

Renewal ROI trends downwards as *P* approaches 1 (Fig. 1D). In each case illustrated here, the lowest *P* values equated to the highest renewal ROI. Biomolecules that initially degrade very rapidly but then transition to a much slower rate of degradation (e.g. mRFP1, mTangerine) exhibit a more dramatic increase in renewal ROI with decreasing *P* than biomolecules with more linear degradation rates (e.g. eGFP). Biomolecules that are considerably more stable at higher *P* values may exhibit a peak in their P-ROI curve, although ROI will always decline as *P* approaches 1 (see Supp. Fig. 1 for a hypothetical example).

### 2.11 Effective Damage Recognition Mechanisms Increase Renewal ROI

To demonstrate the effects of biomolecular damage-recognition mechanisms, renewal selection was next biased towards biomolecules exhibiting the lowest equivalent *P*. The parameter *m* specifies the lowest performing fraction of the population from which the renewal machinery samples randomly upon each renewal event, ranked in terms of equivalent performance. For example, an *m* of 0.30 indicates that those biomolecules with an equivalent *P* in the lowest 30% of all biomolecules constitute the renewal pool of molecules while the remaining 70% of the population (the better performing molecules) will not be selected for renewal. Since biomolecules degrade with time and renewed biomolecules begin life at a high equivalent *P*, a molecule can move between the two pools over time.

By renewing the most degraded biomolecules, a typical renewal event will produce a larger amount of negative entropy compared to a random renewal scenario. With damage recognition, a given *P* can be maintained with a lower rate of renewal (Fig. 2A). In addition, renewal ROI will increase across the entire range of *P*, as demonstrated for all three fluorescent biomolecules (Fig. 2B).

**Fig. 2.**
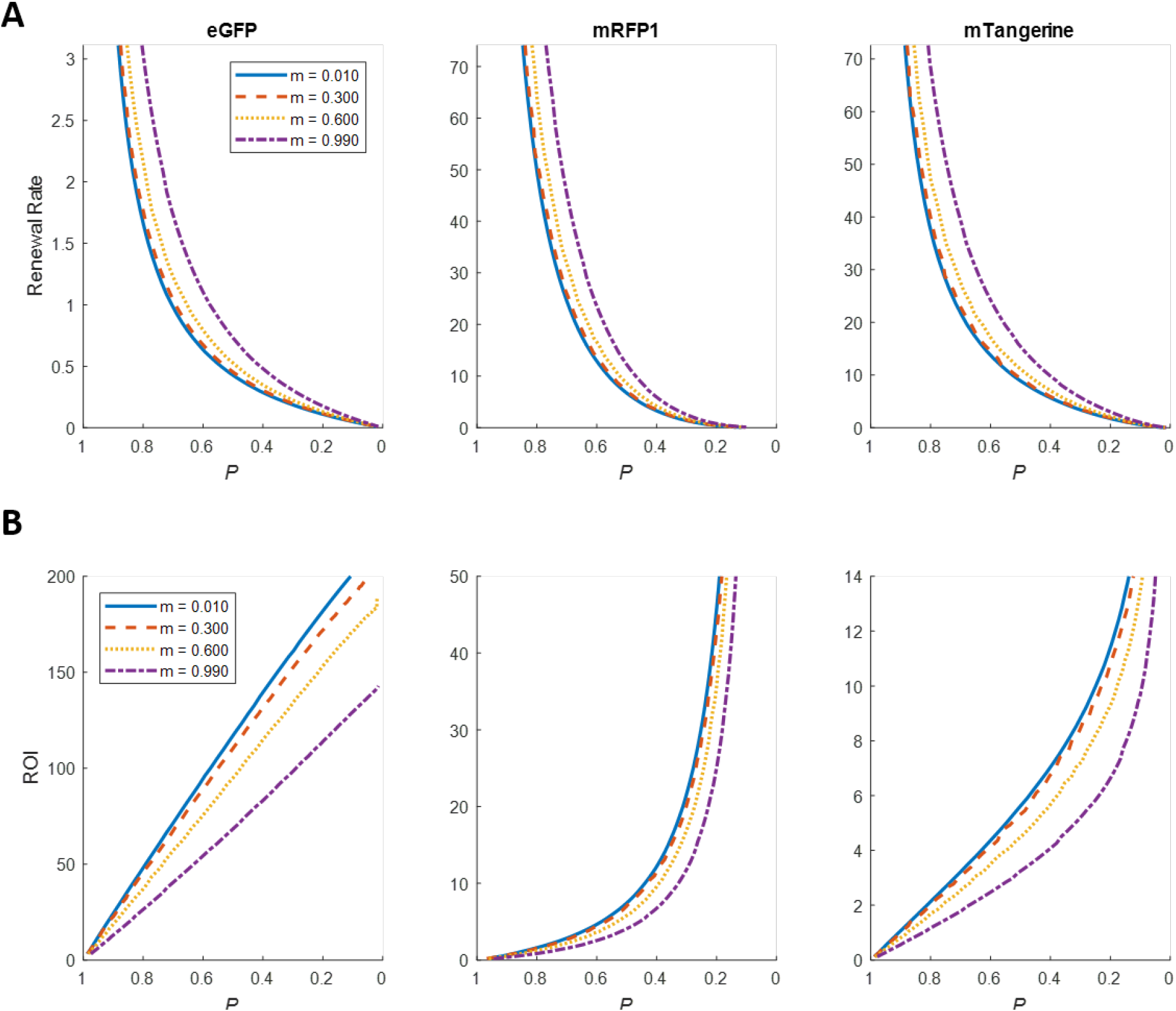
The effect of varying simulation parameter'm' on (a) renewal rate and (b) renewal ROI for eGFP, mRFP1 and mTangerine.

Although renewal ROI improves with damage-recognition ability, the payoffs diminish considerably as *m* approaches zero. For example, the renewal ROI with eGFP at *P* = 0.80 was 78.66% greater than random renewal when *m* was 0.20 (Table 1). Decreasing *m* by a factor of twenty, to 0.01, improved the renewal ROI only marginally compared to *m =* 0.20 (82.30% versus 78.66%). At the other end of the spectrum, a 19.64% increase in renewal ROI was realized with an *m* of only 0.80. These patterns were consistent across all three biomolecules analyzed. This suggests that organisms do not need to invest in extremely precise damage-recognition strategies to receive tangible benefits from damage-recognition capabilities; and furthermore, that extreme precision may not be worth the associated extra costs.

**Table 1.**
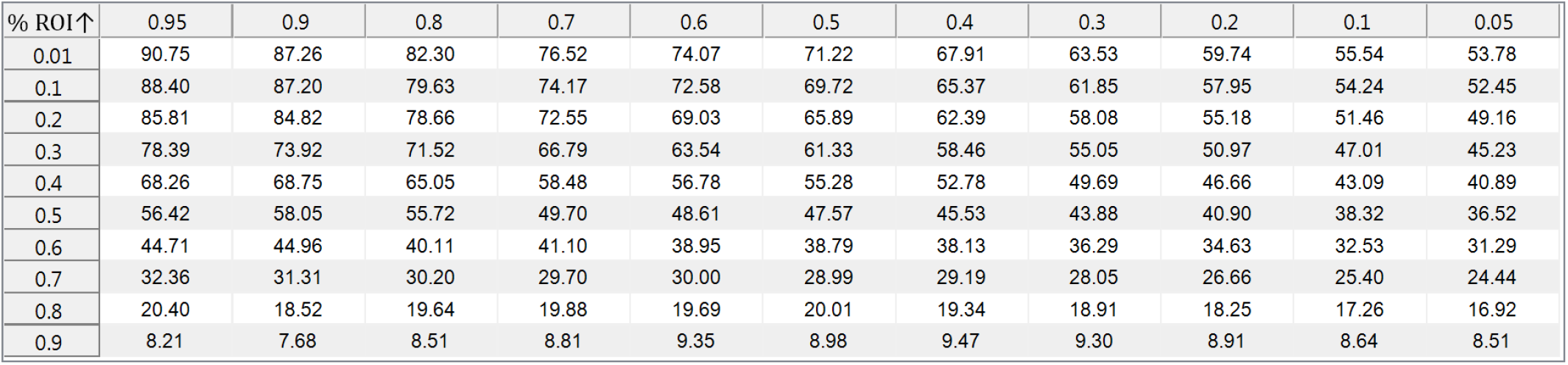
Improvement in renewal ROI for different values of'm' across a range ofP values

*Table values indicate the percent increase in renewal ROI compared to random renewal with eGFP*

## 3 Managing Degradative Internal Entropy Production within an Organism

### 3.1 Extrapolation of Degradation State Concepts to Larger Physical Scales

Although the outlined model was described for a population of biomolecules, a similar thermodynamics model could be applied across a wide range of physical scales. For example, organelles are renewed and face damage due to degradative internal entropy production in a conceptually similar fashion as do the individual biomolecules from which they are assembled. Mitotic cell populations may also be thought of in this way, with individual cells regarded as the renewed unit.

### 3.2 Structural Optimization

Many of the biological mechanisms utilized for producing negative entropy have been characterized. These include biomolecular expression systems, molecular chaperones, degradation systems (proteasomes, lysosomes) and DNA repair enzymes. At the cellular level, stem cells and mitotic cell division, together with apoptosis, provide a means to replace entire cells and, in some organisms, even tissues—thereby maintaining *P* in these populations for a period of time.

Reducing the rate of internal entropy production decreases the amount of negative entropy needed and, in turn, the energetic investment required to maintain homeostasis. The rate of internal entropy production is proportional to the sum of the contributions from all thermodynamic potentials acting on a biomolecular ensemble. This includes chemical reactions, heat, mass, momentum transfer, and other effects. The magnitudes of the thermodynamic potentials depends on the strengths of the respective “damage-inflicting” forces (which may vary significantly with time, particularly when a biomolecule is in an active state) and the ability of an organism's biomolecular structures to resist those forces.

Biomolecular structural optimizations can modulate the effects of degradative thermodynamic potentials by resisting atomic rearrangements. Consider how hydroxyl radicals, which are capable of generating very strong chemical reaction potentials, might affect a protein. The amino acids cysteine and methionine are particularly vulnerable to oxidation reactions (Suto et al. 2006). Substituting another amino acid in place of a cysteine may help to protect a protein from aberrant structural modifications resulting from hydroxyl radical reactions. Alternatively, cysteine could be implemented in non-critical locations within a protein as a sacrificial means to scavenge free radicals and prevent damage to more critical domains. It should be considered, however, that a cysteine or methionine residue in a particular location could bestow an advantageous trait to a protein (improved catalytic activity, energy utilization, substrate specificity, etc.) resulting in an increase in the biomolecule's *Ẇ_m,max_*. Thus, any benefits to inclusion must be weighed against the costs associated with the increased susceptibility to insult.

Other biomolecular structural optimizations that serve to reduce *S_t_* include modifications that improve resistance to undesirable hydrophobic interactions, temperature-induced denaturation, and other alterations. A likely example of these optimizations can be found in the naked mole-rat, whose proteins are extremely resistant to urea-induced unfolding (Pérez et al. 2009).

Biomolecular modifications that optimize stability could result in a deleterious increase in the physical size of the biomolecule (limiting volumetric density) or be otherwise disadvantageous, such as by limiting the realizable rate of intrinsic biological function *(Ẇ_m,max_*). Modifications to biomolecular structure could also affect the amount of resources required for the production or renewal of a biomolecule.

On the other hand, structural alterations that maximize *Ẇ_m,max_* may reduce biomolecular stability and increase *Ṡ_t_*. The polyunsaturated fatty acid content levels of membrane lipids, which varies across species, may represent the evolved configuration that balances biological power density and *S_t_* according the to the specific requirements of a species. This is discussed in section 8.5.

### 3.3 Microenvironment Optimization

The magnitude of the degradative thermodynamic forces acting upon biomolecules is largely determined by microenvironmental conditions. Temperature has a significant effect on reaction velocities and bond forces/energies. Although biomolecular conformation can change with temperature, lower temperature will generally improve molecular stability and reduce *Śļ*. On the other hand, higher temperatures will produce increased kinetic energy transfer during intermolecular collisions, which will increase *Ẇ_m,max_*.

Other attributes of a microenvironment may have less obvious (and sometimes counterintuitive) ramifications. For example, conditions of higher oxidative stress will likely increase the magnitude of degradative thermodynamic forces. At first glance, this may seem purely undesirable from a biological standpoint. Yet, it should be considered that in situations where renewal ROI is highly prioritized such an increase may be acceptable if the ROI benefit from the increased *D* sufficiently overwhelms the downsides of an increase in *Ṡ_t_* resulting from higher oxidative stress. Thus, in situations where an organism has evolved to place extreme priority on renewal ROI (for example, due to limited availability of an energetic resource) and *P* is not as critical a factor, it should not be surprising if the realized *P* is lower in these species. This will present as increased levels of degradation. Conversely, organisms that place lower priority on renewal ROI are likely to function at a higher *P* and thus exhibit lower levels of degradation.

Clearly, optimizing biomolecular structure and microenvironment are multifactorial compromises. Through evolution, relevant parameters are “tested” iteratively and genotypes converge towards those that balance these factors in a way that maximizes species fitness.

### 3.4 The Naked Mole-rat Paradox - Part I

Although oxidative damage is only one of many forms of biomolecular insult that can occur within a living organism, oxidative damage levels are readily measurable and empirically derived values for various organisms have been published. The oxidative damage patterns observed in the naked mole-rat (*Heterocephalus glaber*) are particularly interesting as they are not well explained by any existing aging theory. The naked mole-rat has a maximum lifespan (MLSP) of ˜31 years while its similar-sized cousin, the house mouse (*Mus musculus*), has an MLSP of only ˜4 years (Tacutu et al. 2012). Young naked mole-rats have significantly higher levels of oxidative damage to proteins compared to physiologically age-matched mice (Andziak et al. 2006; Pérez et al. 2009). Furthermore, there is nothing about the naked mole-rat's antioxidant assemblage that would suggest that their antioxidant defenses are any more effective or efficient than that of mice (Andziak et al. 2005).

Naked mole-rats live in a hypoxic environment and have extremely low metabolic rates for their size (McNab 1966). It is considered a paradox that the young naked mole-rat exhibits such high levels of oxidative damage while having extreme longevity compared to similarly sized, closely related species. In fact, this may be predictable and straightforward to explain. The first part of the puzzle is explaining why oxidative damage levels are substantially elevated in the naked mole-rat. The naked mole-rat's limited access to oxygen restricts the rate of cellular ATP production via oxidative phosphorylation, thereby requiring that renewal ROI for cellular processes be highly prioritized. As demonstrated earlier, the steady state conditions that maximize renewal ROI are likely to correspond with increased levels of degradation. Related species that are not as energetic resource-restricted, for example the house mouse, may function at higher *Ps* (to help maximize athletic ability, growth rate, etc.) which will correspond to lower levels of oxidative (and other) damage—at least during youth. Therefore, the elevated levels of oxidative damage found in the young naked mole-rat is likely indicative of the steady state balance between degradation and renewal that maximizes species fitness within the hypoxic constraints of their environment. The second part of solving the naked mole-rat paradox is explaining why their high biomolecular degradation states do not adversely affect MLSP. A possible explanation for this is detailed in section 8.7.

## 4 Thermodynamic Insights into Biological Aging

Youthful organisms are often perceived as existing in a state where biological structure is optimal and essentially perfect. Yet, it is not possible to achieve perfect fidelity in any biomolecular ensemble—all biological systems will exhibit some level of damage, even in youth. In addition, the functional state of biomolecules should not be viewed in binary terms “perfect” and “nonfunctional”, since there are many degrees of performance between these extremes. Rather, it is more accurate to view biomolecular pools as consisting of a distribution of biomolecules across a range of performance/degradation states. As a biomolecule sustains damage, its ability to perform its intrinsic biological function is compromised—but this will likely not manifest as an all or nothing scenario. For example, biomolecular fluorescence decreases over time as the molecules sustain (predominantly oxidative) damage during the course of doing work but, typically, many separate insults are required to completely eliminate the capacity for fluorescence emission.

### 4.1 Energetic Expenditures towards Biomolecular Repair and Replacement - a Paradox?

Proteins account for the majority of biomolecules within a cell. The rate of total protein synthesis has been empirically determined for a number of species (Table 2). Smaller organisms synthesize proteins at higher rates than larger species (Fig. 3A). In mice, protein synthesis takes place at a rate sufficient to replace total body protein mass every ˜5.3 days, while a human requires ˜39.3 days. Of course, renewal rates for individual proteins can vary widely protein-to-protein, ranging from minutes to years (Hetzer 2013).

**Fig. 3.**
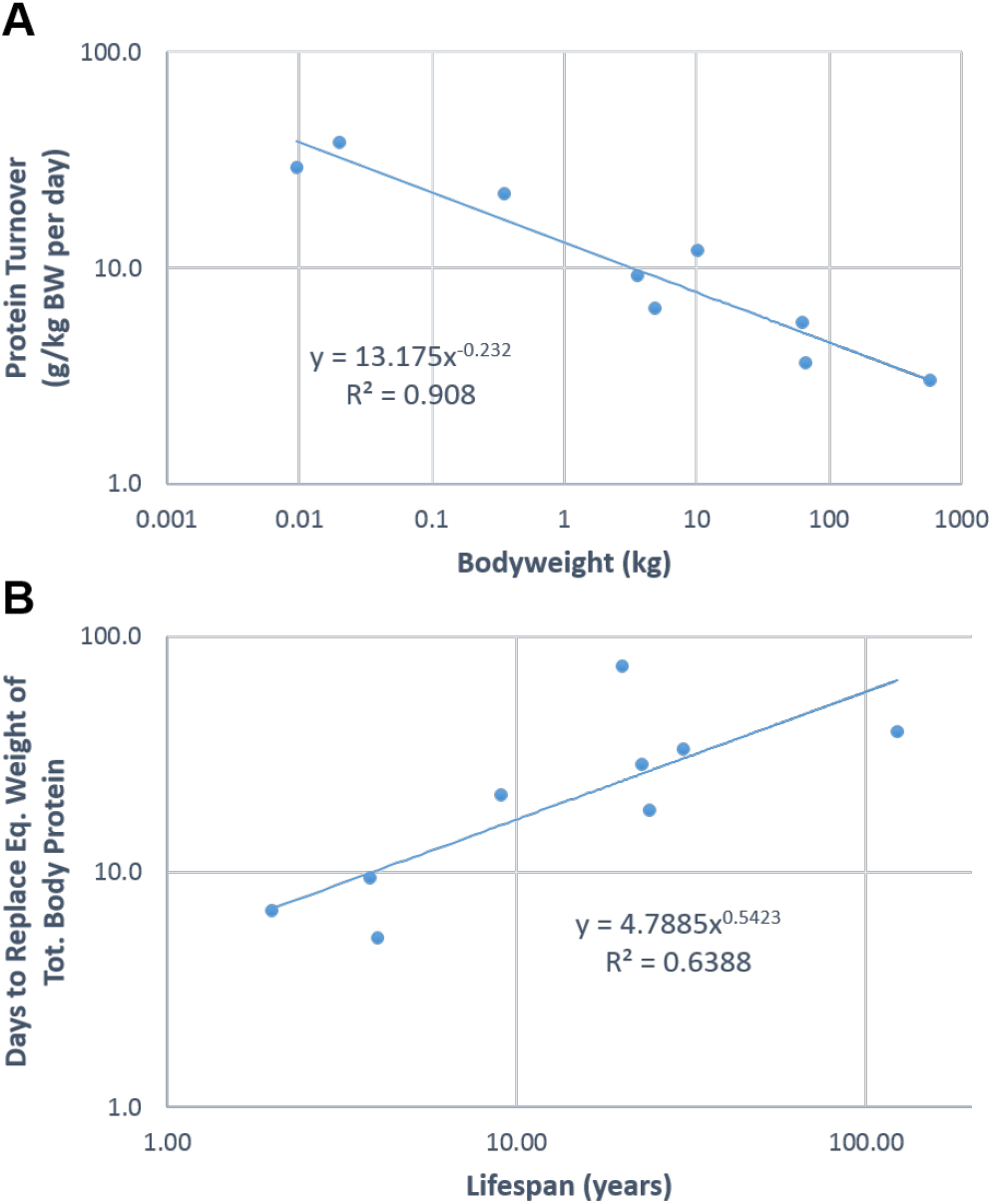
(a) Protein turnover as a function of bodyweight for the species listed in Table 2 and (b) days to replace the equivalent weight of total body protein as a function of MLSP. Data from Table 1.

**Table 2.** Protein Synthesis Rates, Number of Days to Renew Total Body Protein Mass, and Number of Renewals per Lifespan for Different Metazoan Species

| Species | BW (kg) | Protein synthesis |  |  | Body Protein Composition (%) | Days to Renew Total Body Protein Mass | Maximum Lifespan (years)<sup>10</sup> | Renewal Per Life |
| --- | --- | --- | --- | --- | --- | --- | --- | --- |
|  |  | g/day | g/kg BW per day | g/kg<sup>0.75</sup> BW per day |  |  |  |  |
| Honey possum | 0.010 | 0.280<sup>1</sup> | 29.1 | 9.1 | (17.0)<sup>†</sup> | 6.9 | 2.0 | 106 |
| Mouse, small | 0.020 | 0.768<sup>2</sup> | 38.4 | 14.4 | 20.3<sup>7</sup> | 5.3 | 4.0 | 277 |
| Rat | 0.35 | 7.7<sup>3</sup> | 22.0 | 16.9 | 20.8<sup>7</sup> | 9.5 | 3.8 | 147 |
| Rabbit | 3.6 | 33.1<sup>3</sup> | 9.2 | 12.7 | 19.4<sup>7</sup> | 21.1 | 9.0 | 155 |
| Cat (HP) | 4.8 | 31.4<sup>4</sup> | 6.5 | 9.7 | 21.8<sup>7</sup> | 33.3 | 30.0 | 328 |
| Dog | 10.2 | 123.4<sup>5</sup> | 12.1 | 21.6 | 22.1<sup>7</sup> | 18.2 | 24.0 | 480 |
| Sheep | 63 | 351<sup>3</sup> | 5.6 | 15.7 | 16.0<sup>8</sup> | 28.7 | 22.8 | 290 |
| Man | 67 | 245<sup>6</sup> | 3.7 | 10.5 | 14.4<sup>9</sup> | 39.3 | 122.5 | 1137 |
| Cow | 575 | 1740<sup>3</sup> | 3.0 | 14.8 | 22.5<sup>7</sup> | 74.4 | 20.0 | 98 |

<sup>†</sup>Body protein concentration for honey possum not available, used 17.0% for “days to renew” calculation. <sup>1</sup>Bradshaw and Bradshaw, 2009. <sup>2</sup>Garlick and Marshall, 1972. <sup>3</sup>Reeds and Harris, 1981. <sup>4</sup>Russell et al., 2003. <sup>5</sup>Everett et al., 1977. <sup>6</sup>Pacy et al., 1994. <sup>7</sup>Moulton, 1923. <sup>8</sup>Reid et al., 1968. <sup>9</sup>Mitchell et al., 1945. <sup>10</sup>Tacuta et al., 2012.

Over the course of a lifetime, a long-living human will synthesize enough protein to replace total body protein mass 1137 times over (Table 2). Let us assume for the moment that degraded/dysfunctional proteins are “accumulating” as an individual grows older due to insufficient energy expenditure toward repair and replacement, as suggested by the disposable soma theory of aging (Kirkwood 1977; Kirkwood and Holliday 1979; Kirkwood and Rose 1991). Considering a worst-case scenario where all protein in an aged human is in need of replacement, it would only require an estimated 0.09% increase in daily resource investment in protein synthesis to offset the average daily increase in degraded protein. This translates to 0.23 calories per day^4^. With a daily dietary intake of 2500 calories, this is a mere 0.0092% of daily energy intake. Protein synthesis represents the majority of the total energy spent on biomolecular renewal, and the total amount of protein dedicated to translation is 2-15 times greater than that dedicated to transcription and DNA maintenance (Liebermeister et al. 2014). In light of the very small additional investment needed to offset an increase in protein degradation, the disposable soma theory's claim that aging is caused by an energetic underinvestment in repair and maintenance that results in an accumulation of damage (Kirkwood 1977; Kirkwood and Holliday 1979; Kirkwood and Rose 1991) is difficult to accept. In addition, organisms that renew their proteins more frequently generally exhibit decreased longevity—not increased (Fig. 3B). Smaller organisms renew protein at a faster rate than larger organisms (Fig. 3A) but have reduced longevity (Calder 1984; Austad 2005; de Magalhães et al. 2007).

A corollary of the disposable soma theory suggests that an increase in available energy would allow an organism to devote more resources to somatic maintenance and thus extend longevity. However, no studies exist demonstrating that increased caloric intake extends longevity—while it is well known that obesity leads to decreased longevity and diabetes (Ahima 2009). It is clear that energetic expenditures towards renewal alone cannot explain the differences in longevity between species nor does it provide a solid rationale to explain why aging must occur in the first place.

### 4.2 Biomolecular Degradation and Aging - Accumulation versus Homeostatic Shifts

Aging has been described as the accumulation of unrepaired damage. This implies that the biomolecular degradation in an aged individual results from lifelong accumulation. Perhaps this assessment is not entirely accurate.

Organisms exhibit biomolecular degradation even during youthful homeostasis. Reducing the rate of biomolecular renewal from the youthful level (or any other level) will decrease *P* and increase the perceivable amount of degradation. Once steady state is reestablished around a new renewal rate, a phenomenon that could be described as a “homeostatic shift”, no further reductions in *P* should occur—unless outside factors are at play.

Restoration of proteasome function in aged human dermal primary fibroblasts largely restores markers of protein aging to youthful levels (Hwang et al. 2007). This is analogous to a shift in the protein pool from a lower to a higher *P*, and demonstrates that the reduction in *P* occurring with age may be at least partially reversible. If the degraded protein was truly representative of accumulated, irreparable damage then upregulation of proteasomal function should not have eliminated the damage. The fact that the condition is essentially reversible suggests that the increased biomolecular degradation found in aged individuals is not due to damage “accumulation” but is more likely attributable to reduced biomolecular renewal leading to a corresponding reduction in *P* and thereby an increased degradation state^5^. Consistent with this notion, protein renewal does indeed significantly decline during aging (Richardson and Cheung 1982; Rattan 1996; Ryazanov and Nefsky 2001).

A distinction between *accumulated* dysfunctional biomolecules and a shift in biomolecular degradation state caused by reduced renewal can be made by determining whether renewal is actively occurring. Damage in molecules that are eventually renewed should not be referred to as “accumulated” damage, even if the rate of renewal has decreased and the degradation state of the biomolecular pool is high.

This raises further doubt over the disposable soma theory's assertion that aging is caused by an energetic underinvestment in repair and maintenance resulting in an accumulation of damage (Kirkwood 1977; Kirkwood and Holliday 1979; Kirkwood and Rose 1991). The idea of an “energetic underinvestment” is a misnomer, as no amount of energetic investment will produce a perfect population of biomolecules (i.e. a degradation state of zero). Increasing biomolecular renewal will reduce *D* but renewal ROI will continually worsen as renewal rate is increased. An energetic underinvestment in renewal cannot explain why youthful homeostasis is lost—there must be a higher-level initiating event. *Declining energetic expenditures towards renewal are merely another consequence of aging, not a root cause*. This begs the question of why renewal rates drop with age in the first place.

### 4.3 Total Entropy Increases Slowly with Age in Comparison to Internal Entropy Production Rate

As highly ordered entities existing in thermodynamic nonequilibrium, living organisms are subject to internal entropy production leading to ongoing molecular damage. The high frequency at which the animals depicted in Table 2 replace their total body protein mass demonstrates that the rate of degradative internal entropy production *Ṡ_i,ind_* within an individual organism is much greater than the rate at which the total structural entropy of the individual increases with age post reproductive maturity *Ṡ_tot,ind_*.

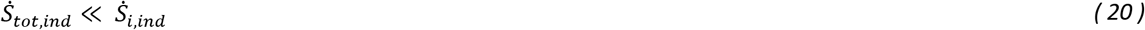

Eq. *(20)* is intuitively evident when the speed at which biological material degrades at biologically relevant temperatures, even when conditions are sterile and optimized, is contrasted against the period of time that organisms—including those with only moderate longevity—are alive.

Cellular mechanisms work towards counteracting damage resulting from internal entropy production, coming close to establishing a steady state in terms of preservation of biomolecular integrity when observed over a limited time window. Despite the efforts of these renewal systems, the overall degradation state (i.e. the total structural entropy) of an individual organism increases with age post reproductive maturity. The aging process could be viewed as a progression through many discrete nonequilibrium steady states that eventually result in a global degradation state that renders the individual nonviable. Why then do organisms transition between these states and why is youthful homeostasis lost? The answer to this question may lie in how an organism combats internal entropy production affecting nonrenewable structures, i.e. those subject to irreversible losses in fidelity.

## 5 Preservation of DNA Molecular Information

Most types of biomolecules are replaced by expression of a genetic sequence or are the metabolic products of expressed biomolecules. The performance of these biomolecular pools can be preserved by the successful expression of the appropriate genetic sequence(s) or the relevant metabolic processes, and the removal or repair of any dysfunctional counterparts. With a given rate of renewal and assuming intact expression machinery, the preservation of biomolecular performance within a cell becomes dependent on: (1) the fidelity of the genetic material responsible for biomolecular expression, and (2) the cell's ability to remove all dysfunctional biomolecules. While the last requirement should not be trivialized, this is a very attainable objective: that is to say, the specificity of degradation pathways can afford to err on the side of casting a wider net to help ensure that any dysfunctional biomolecule is eventually recognized since these biomolecules can be resynthesized anew. Indiscriminate purging of cellular content would eventually dispose of any unwanted products. For these reasons, fidelity preservation in biomolecules that are not expressed, the genetic-encoding biomolecules, particularly warrants further scrutiny.

### 5.1 DNA Informational Fidelity Loss is Unavoidable

DNA molecules contain the information encoding for production of all other classes of biomolecules as well as the orchestration of all cellular processes. They are unique among classes of biomolecules as they depend on their own integrity for replacement. Like all molecules, DNA is subject to damage resulting from internal entropy production and will display an insult rate proportional to the damage-inflicting thermodynamic potentials of its microenvironment.

There are a number of ways that DNA damage can occur, resulting in base alterations, cross-linking, strand breaks, and other modifications (De Bont and van Larebeke 2004). For example, consider some of the possible outcomes after a double-stranded DNA molecule has suffered a single base excision:

1. The damage is properly repaired by endogenous base excision repair (BER) mechanisms
2. The damage is improperly repaired by BER mechanisms
3. Additional insults occur at the damage site before repair can take place
4. No repair or further damage occurs for a length of time

DNA replication takes place far from thermodynamic equilibrium. The accuracy of DNA polymerase is largely dependent on the differences in the free energy released or absorbed by the various possible base-pairing combinations of incoming nucleotides to template strand nucleotides (Arias-Gonzalez 2012). Utilizing thermodynamic theory, researchers have estimated polymerase error rates and have shown them to be non-zero (in alignment with empirical findings). Although BER often successfully repairs single-base excision damage (scenario 1)—restoring redundancy and preventing changes in stored information—there is always a possibility that a replication error will occur. Additionally, repair machinery must translocate to the site of the insult and perform the repair. This will not occur instantaneously. If further damage occurs at the site before repair takes place then information could be permanently lost.

A single level of redundancy is definite at all DNA base pairs—that provided by the pairing base on the opposite strand. Even an insult restricted to a single base will deplete this redundancy and can lead to a permanent change in DNA information. This does not imply that insults that are more serious are not repairable. For example, double-stranded breaks can be repaired by homologous recombination in many cases, but there is no guarantee that a homologous site exists or that the repair will be successful.

Once a DNA molecule has suffered an insult, there is no means to ensure restoration of redundancy. Sporadic reversions are possible but by no means inevitable. *As the second law stipulates that molecular insults are inevitable, the genetic data stored in DNA molecules must change with time—indefinite preservation of data is not possible. The concept of “perfect” DNA repair is flawed and unattainable*. This same inference has been drawn previously utilizing information theory: Yockey (1974) suggested that the noisy-channel coding theorem stipulates that, under ideal conditions, the stability of the genetic message can be such that the error is “arbitrarily small” but that the probability of error must always be non-zero.

Permanent losses in genetic fidelity are typically discussed in terms of discrete mutations or mutation rate (Sniegowski et al. 2000). Perhaps a better way to assess these losses is to utilize the concept of mutual information^6^, which is a measure of the amount of information shared between two datasets. This provides a means to quantitate the amount of encoded data retained in, or transmitted between, DNA molecules over time. As genetic data must change with time, the mutual information of a discrete, non-replicating DNA molecule will decrease with time.

DNA insults do not only affect mutual information, they can also result in an increase (or decrease) of total information. For example, insertions, duplications, and repeat expansion mutations increase the total amount of information. Although mutual information will always trend downwards, it is important to recognize that the total amount of DNA information could increase, decrease, or stay the same over time.

A means to quantify DNA informational fidelity *Φ_DNA_* is to consider the ratio of mutual information shared between the two states being compared *I_mutual_* to the maximum of the total amount of information in the initial *I_init_* and final *I_final_* states.

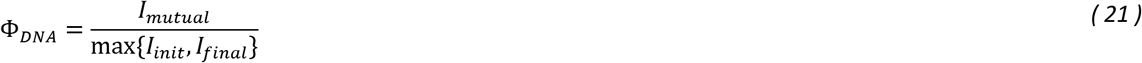

Only in the case where no original information has been lost and no new information gained will Φ_DNA_ be equal to its maximum possible value of 1.

### 5.2 Applicability of the Degradation State Concept to DNA Molecular Ensembles

Synthesized biomolecules depend on the fidelity of DNA for their correct expression. If the full fidelity of DNA is preserved then, theoretically, negative entropy could be produced at a rate that maintains a steady *P* in any expressed biomolecular pool. Since DNA molecules rely on their own fidelity for identical replacement, this biomolecular replacement scenario is not applicable to DNA molecules.

### 5.3 Considerations for DNA Informational Fidelity Preservation in Different Cell Types

For any sexually reproducing multicellular organism, the zygote contains the truest representation of the parentally derived genetic data anywhere in the individual and of any stage in life, i.e. the DNA informational fidelity between parent and offspring is maximal in the zygote. The informational fidelity of an organism's DNA at any later point in life can be quantified by comparing to this baseline standard.

Consider how the requirements for preservation of DNA informational fidelity are likely to vary over the course of an individual multicellular organism's life and as a function of cell type. Somatic cellular function must remain at a sufficiently high level for some minimum amount of time to allow the organism to reproduce. Selective pressure for preservation of function begins to decrease as an individual ages past reproductive maturity (Medawar 1952; Hamilton 1966). The progeny of adult stem cells are the replenishment source for somatic cells; therefore, the DNA informational fidelity of adult stem cells must be greater, on average, than non-stem somatic cells for any given point in an individual's life.

The redundancy provided by diploidy/polyploidy, gene duplication, and functional overlap provides a degree of robustness that enables non-germ cells to tolerate some amount of DNA informational fidelity loss with minimal performance impact on the individual (Medvedev 1972; Yockey 1974; Riggs 1994; Plata and Vitkup 2013). Similar levels of damage would be more detrimental in germ cells as they would propagate to all cells of the progeny. Therefore, the average DNA informational fidelity of germ cells must be greater than that of adult stem cells at the time of reproduction, which in turn must be greater than the DNA informational fidelity of nonstem somatic cells at the time of reproduction. This relation can be written

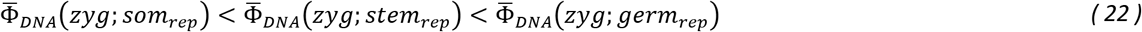

where 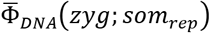 represents the average DNA informational fidelity of non-stem somatic cells of an individual at the time of reproduction referenced to the same individual when it was a zygote, 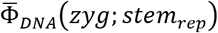 is the average DNA informational fidelity of adult stem cells referenced to the zygote, and 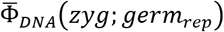 is the average DNA informational fidelity of germ cells referenced to the zygote.

In agreement with Eq. (22), organisms appear to come closest to fully preserving DNA informational fidelity in germ cells. This results from evolved strategies that place extraordinary emphasis on the preservation of both nuclear and mitochondrial genetic data in germ cells. The fidelity of mtDNA is largely restored during oogenesis through a genetic bottlenecking process that selects for the healthiest mtDNA and eliminates mutated mtDNA molecules that compromise mitochondrial function (Wai et al. 2008; Lee et al. 2012). The nuclear DNA in germ cells is subject to very strict insult detection mechanisms (Hochwagen and Amon 2006; Jaramillo-Lambert et al. 2010; Bailly and Gartner 2013). Germ cells are more likely than somatic cells to undergo apoptosis when DNA damage is detected, rather than attempt to repair the damage (which often results in the loss of DNA informational fidelity). They are also sequestered in a protected microenvironment with various specialized cells whose sole purpose is to support and maintain the germ cells (Schulz et al. 2002).

Assessing the situation from a thermodynamics perspective suggests that the rate of DNA informational fidelity loss is minimized by keeping the thermodynamic potentials acting on DNA molecules as low as possible to reduce the rate of molecular insult. Primordial germ cells (gametogonia), as well as oocytes and spermatocytes, have relatively low rates of oxygen consumption (Brinster and Troike 1979). Most adult stem cells are quiescent and frequently prioritize glycolysis over oxidative phosphorylation, leading to lower levels of free radicals and reactive oxygen species (ROS) (Tothova et al. 2007; Rossi et al. 2008; Suda et al. 2011; Shyh-Chang et al. 2013) and decreased mtDNA replication rates. This supports the notion that manipulation of thermodynamic potentials acting on DNA molecules, through modulation of cellular processes and microenvironmental conditions, is a realizable and effective means of reducing the rate of DNA informational fidelity loss in cells.

The genetic similarity between gametes and the same individual when it was a zygotes Φ*_DNA_*(*zyg*; *gametes*) is determined by not only inevitable germ cell DNA informational fidelity losses throughout life but also losses due to genetic recombination during meiosis ΔΦ_recom_:

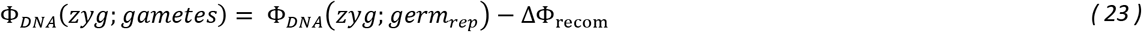

Singular insult events that generate losses in DNA informational fidelity (i.e. mutations) most commonly have little or no effect on offspring fitness. Some mutations will result in decreases in fitness while only the rare insult produces increased fitness (Fisher 1930; Eyre-Walker and Keightley 2007). Absent effective evolutionary selection, genetic dissimilarities (resulting from the loss of mutual DNA information between parent and offspring, and/or potential gain of new information) will lead to a decline in species fitness since advantageous mutations are rare. The proportion of progeny with low fitness must not be so excessive that evolution cannot successfully select for neutral and higher fitness offspring. For this reason, a minimal limit Φ*_DNA,min_*(*zyg*; *gametes*) is effectively placed on the DNA informational fidelity of the progeny as referenced to the parent:

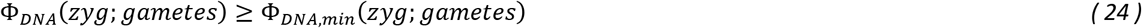

Germ cells must be maintained with adequate redundancy levels and a sufficiently stringent fidelity preservation strategy to satisfy Eq. (24). In this way, DNA informational fidelity is largely—but not perfectly—preserved generation-to-generation and the relatively small fidelity losses that do occur can be eliminated from the gene pool if they negatively affect species fitness, or retained when they increase species fitness.

There is a direct correlation between the lifetime risk of cancer in a tissue and the number of divisions of the stem cells maintaining that tissue (Tomasetti and Vogelstein 2015). The strategies used to preserve stem cell fidelity clearly do not achieve the fidelity preservation of germ cell strategies. Since the preservation of stem cell DNA informational fidelity requires dedicated niches with specialized microenvironments, there must be associated negative fitness costs to scaling these niches excessively—even though doing so may result in further reductions in the rate of DNA informational fidelity loss in the cell type in question. For this reason, an organism's stem cell niches must adequately support the respective target tissue over the lifespan of the individual, but not be so unnecessarily burdensome that they lower species fitness.

## 6 Establishing a Connection between Thermodynamic, Information, and Evolutionary Theory in Biological Aging

Modern nonequilibrium thermodynamic theory stipulates that all biomolecules will suffer degradative insults. Biological repair and replacement mechanisms cannot guarantee that DNA informational fidelity is preserved in individual cells regardless of the level of investment: Cellular DNA informational fidelity must decrease with time. Per Eq. (21), this is sufficient to mandate a decrease in DNA informational fidelity. What are the repercussions of these losses for an individual, species as a whole, and discrete genes within a species genome?

### 6.1 In the Individual

Most germline mutations are neutral or detrimental to fitness, with only the rare mutation being beneficial. It follows that mutations occurring in the somatic cells of an individual organism would exhibit this same pattern with regards to their contribution towards the viability of the individual. Therefore, without selection for only those changes that are neutral or beneficial to the individual, DNA informational fidelity loss in somatic cells will reduce individual viability with time, i.e. individual organisms would age.

Although natural selection is traditionally considered to occur at the level of the individual, a similar process takes place during the life and death cycles of the individual cells of a multicellular organism throughout its life; single cells are the replicating unit in this scenario. For an individual multicellular organism containing cells undergoing mitosis, natural selection will occur on the cellular level and favor those cells that display the highest fitness. These configurations may not necessarily be the most beneficial to the individual as a whole. As natural selection will always be present at the level of the replicating unit (Szathmáry and Smith 1995; Baum et al. 2013)— cells in this case—the individual must rely on imperfect innate biological mechanisms that attempt to select for only those configurations that do not reduce overall organismal viability.

As there is no means for an organism to perform comparative DNA sequence analyses, cells with undesirable base-sequence modifications are only detectable by phenotype. In the case of more severe damage, cellular mechanisms enable detection of the damage and initiation of apoptosis (Zhou and Elledge 2000). On the other hand, singular mutations may exhibit very mild or no detectable undesirable phenotype. For example, mutations whose effects are masked by redundancy are likely to have no detectable phenotype. A mutation may also occur in a region of the genome that is currently inactive or irrelevant to the particular cell—therefore there may be no immediate negative phenotype. Such a genomic region could become active later, at which time the mutation may have already spread to the cell's progeny.

Even in the ideal embodiment, the effects of multiple mutation events must eventually decrease individual viability; at some point, removing cells determined by biological mechanisms to be undesirable will no longer provide reprieve from losses in viability since cellular DNA informational fidelity will continue to decrease until all cells approach the detectable threshold of dysfunction. At this point, there would be no "good" configurations to select for to replace those cells determined to be undesirable, even if such cells could be detected with perfect accuracy.

Genetic redundancies are likely able to temporarily compensate for the loss of DNA informational fidelity in an individual—essentially delaying a detectable aging phenotype to at least the age of reproductive maturity (Fig. 4A). A second line of defense is provided by innate mechanisms that identify specific types of cellular dysfunction and eliminate cells displaying those phenotypes (Zhou and Elledge 2000). Once the utility of these redundancies is expended and ever-increasing numbers of compromised cells circumvent innate detection mechanisms, the individual will no longer be able to avoid a loss in viability. The resulting dysfunction becomes progressively worse with time. *As no existing, or theoretical, biological means has been demonstrated or postulated to be capable of selecting only for those changes in cellular DNA information that are neutral or beneficial to the individual, the second law stipulates that all individual organisms must eventually age if they live long enough*. The claim by Hamilton (1966) that, at the species level, senescence arises inevitably due to declining selection pressure with age, while not challenged here, is redundant.

**Fig. 4.**
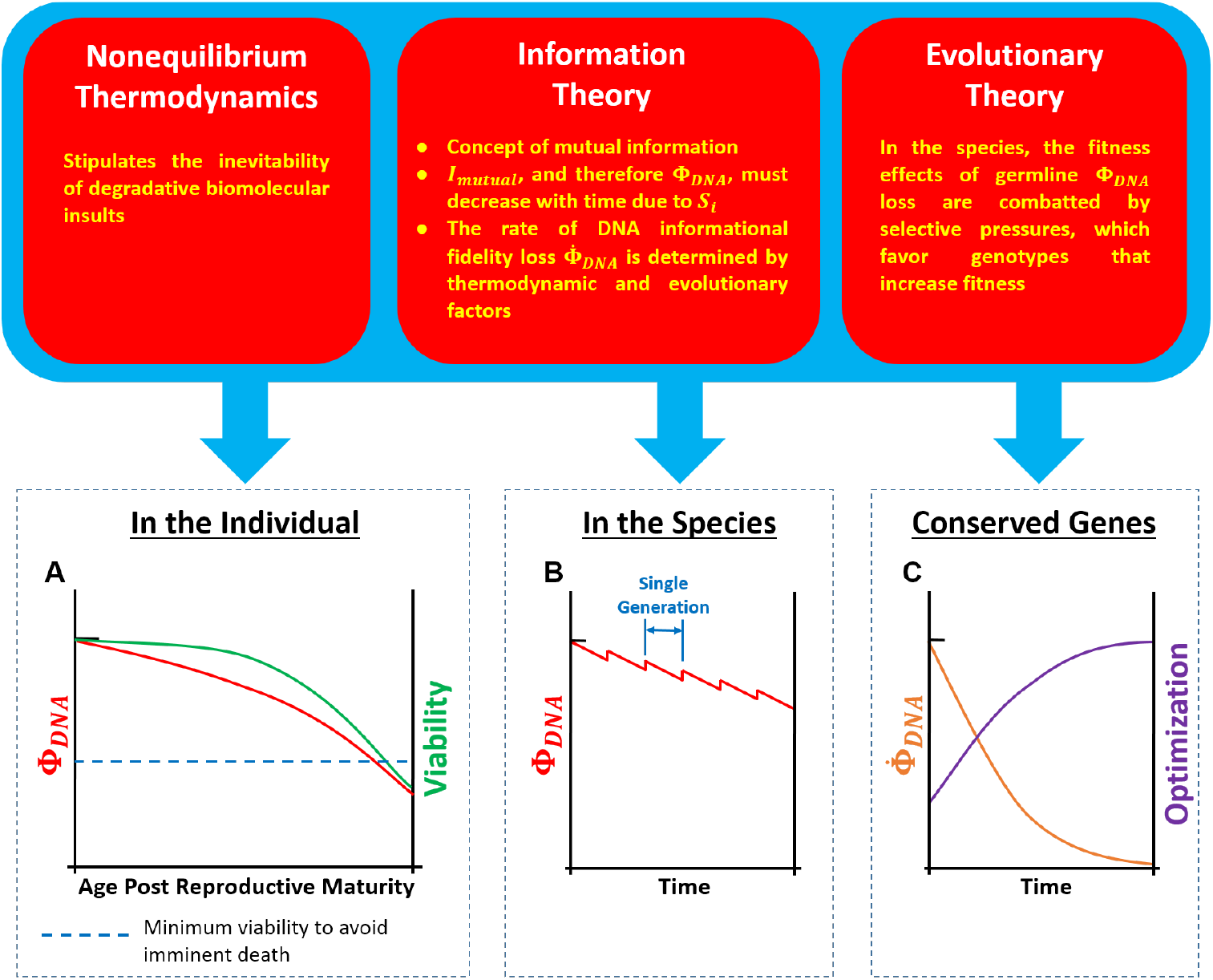
The proposed connection between thermodynamics, information, and evolutionary theory in generating mandatory DNA informational fidelity losses in individual organisms, the genomes of species, and individual genes within a species genome. (a) Although a correlation between individual viability and somatic cell DNA informational fidelity loss is expected, genetic redundancies and other compensating mechanisms may attenuate reductions in viability generated by DNA informational fidelity losses as individuals age. (b) The average DNA informational fidelity in cells of all individuals of a species, as measured against a common ancestor, will decline as the individual ages. This loss can be largely reverted in subsequent generations by sequestering germ cells in conditions optimized for preservation of genetic data. (Generations are aligned for illustration purposes.) (c) In conditions of static selective pressure, such as may be approximated with highly conserved genes, the rate of DNA informational fidelity loss will decrease and approach zero as gene optimization approaches a maximal value.

In addition, since there is no possible means to prevent natural selection from occurring at the level of the individual cell^7^, the second law also stipulates that cancer is inevitable in any individual organism given sufficient time—despite the fact that cancer has yet to be detected in a small number of studied species. The naked mole-rat has long been considered one such species. This was challenged in two recent articles that reported cancer in the naked mole-rat for the first time (Delaney et al. 2016; Taylor et al. 2016).

### 6.2 Within a Species

Due to gamete fidelity preservation mechanisms (Jaramillo-Lambert et al. 2010; Bailly and Gartner 2013), the loss in DNA informational fidelity suffered between a gamete and the zygote that gave rise to the individual that produced the gamete will typically be less than the average loss in somatic cells of the individual (at the age of reproductive maturity). By limiting the DNA informational fidelity loss in the gamete to a low enough level that selection for neutral and higher fitness offspring is possible, species fitness can be preserved and even increase—as can complexity. Natural selection is the only means to prevent inevitable DNA informational fidelity loss from generating mandatory reductions in species fitness. Since DNA insults can result in new information, the second law does not constrain complexity; through selection, complexity can increase as warranted to maximize fitness.

Fluctuations in species selective pressures result in adaptive genetic changes that drive species DNA informational fidelity (as measured relative to a common ancestor) downwards with time (Fig. 4B). Since the process of evolving mandates genomic changes, conditions where selective pressures are changing at a high rate can lead to DNA informational fidelity losses in the species that are greater than the loss predicted by Φ*_DNA_*(*zyg*; *gametes*).

### 6.3 Within Discrete Genes

A number of genes are highly conserved even amongst diverse species whose most recent common ancestor dates back hundreds of millions of years. If DNA informational fidelity loss is inevitable, how can the rate of DNA informational fidelity loss in these genomic sequences be so low?

With highly conserved genes, selective pressures remain very static and common between organisms. There are a number of reasons for this. Firstly, highly conserved genes typically have extremely specific functions that fulfill common needs across the range of species represented. Many of the genes with the most fundamental and specific functionality—e.g. ribosomes, tRNA, histones— also tend to be the most conserved (Isenbarger et al. 2008). Since an individual gene is far less complex than the organism it constitutes, there are considerably fewer ways for a highly conserved gene to achieve increased optimization to perform its intrinsic biological function. Thus with time these types of genomic sequences converge around a common solution that facilitates maximal fitness in a range of species. The second law remains very much active in these cases—i.e. DNA mutations still occur. However, selective pressures have a greater negating effect against DNA informational fidelity losses in these highly conserved genetic sequences.

Consider the evolution of a highly conserved genetic sequence (prior to species divergence) under conditions of static selective pressures. The rate of DNA informational fidelity loss Φ*_DNA_* will assume some initial value (Fig. 4C). Through the course of evolution, fewer configurations will be available that are more optimized than the current configuration. As a result, the rate of DNA informational fidelity loss decreases and the rate of optimization is also reduced. With fewer "positive" mutations available and selection tending to eliminate (i.e. revert) "negative" mutations, DNA informational fidelity losses are minimized. As the optimization of the genetic sequences approaches a theoretical maximum, the rate of DNA informational fidelity loss approaches zero. Essentially, those losses produced as a consequence of the second law are effectively negated by a gain in DNA informational fidelity resulting from selective pressures.

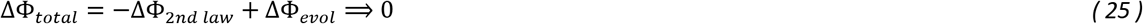

This can only arise when 1) selective pressures are essentially static and 2) a very long period of time has passed, and tends to obfuscate the stipulated effects of the second law on highly conserved genetic sequences.

Compared to individual genes, many more factors come into play in defining selective pressures for an entire species. As it is incredibly unlikely for all of these factors to remain stable for any length of time, conditions of perfectly static species selective pressure are not realizable and the genomes of species, considered as a whole, do not approach this theoretical lower limit of DNA informational fidelity loss. Therefore, the ramifications of the second law are more easily discerned when examining DNA informational fidelity losses in the full genomes of species over time.

## 7 Entropy-Driven Managed Deterioration - Basic Concepts

Having identified one class of biological structure universally subject to mandatory and irreversible losses in fidelity, we return to our previous question of why organisms undergo homeostatic shifts as they age.

Fig. 5 depicts the basic interrelationships that may explain the progression of the aging phenotype in many metazoans. In this model, the key top-level factor initiating the transition from youthful homeostasis is internal entropy production, which inevitably generates losses in DNA informational fidelity in both mitochondrial and nuclear DNA. Informational fidelity losses in mitochondrial DNA (mtDNA) are ubiquitous in aged mammals (Wallace 1999) and lead to lower peak energy output (Yaniv et al. 2013). This decline in mtDNA informational fidelity is partially modulated by a controlled deceleration in mitochondrial biogenesis (Figge et al. 2012), which reduces the rate of clonal expansion of degraded mtDNA and limits the exposure of mtDNA to the high thermodynamic stress conditions of replication events. The escalating deficit in cellular energy currency (ATP) production gives rise to a progressively worsening inability to fund all cellular processes at youthful levels, and generates forced reductions in biomolecular turnover that increases biomolecular degradation state and lowers biomolecular performance—representative of a transition away from youthful homeostasis.

**Fig. 5.**
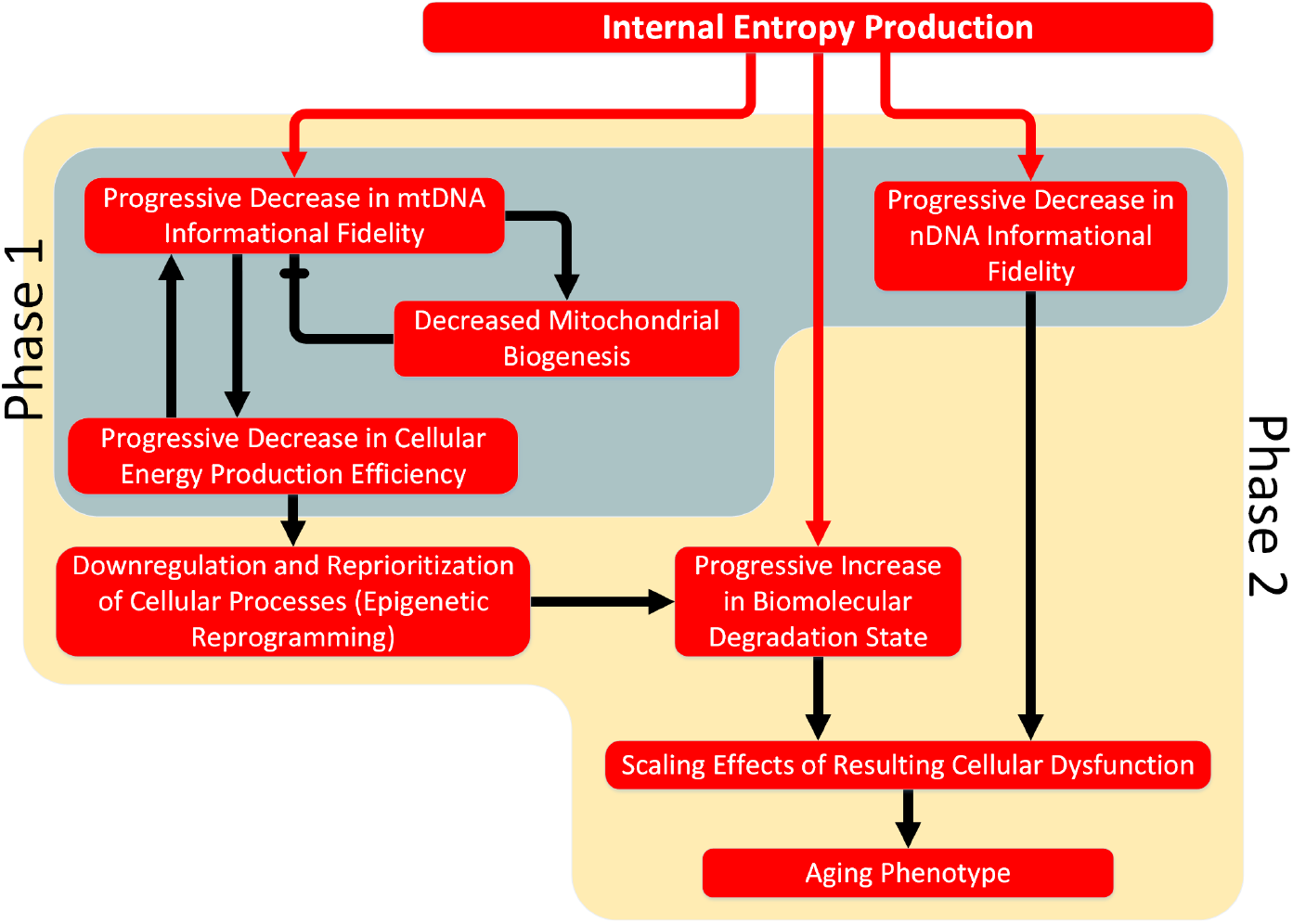
The basic interrelationships between primary factors that may largely describe the progression of the aging phenotype in many metazoa. During 'Phase 1' of an individual's life, DNA informational fidelity losses have not reached levels sufficient to generate an aging phenotype. *'Phase 2'* begins when dysfunction has progressed to the point that aspects of the aging phenotype begin to become apparent.

Concurrent with these changes, increasing losses in nuclear DNA fidelity with age (Podolskiy et al. 2016) produce a mosaic of stochastic cellular dysfunction that worsens over time (Bahar et al. 2006; Lodato et al. 2015). Together with the described mitochondrial dysfunction, this could largely explain age-linked cellular dysfunction and the overall aging phenotype of the organism. "Longevity optimization" genes may have evolved to attenuate the negative effects of informational fidelity losses in nuclear and mitochondrial DNA through reallocation of resources and physiological alterations. This model is discussed in more detail in section 9.

## 8 Longevity Determination

Leonard Hayflick has stated that aging is not driven by genes but by thermodynamics (Hayflick 2004), while also arguing that the genome does govern longevity. Additionally, Hayflick maintains that natural selection has led towards biomolecular arrangements that are capable of preserving fidelity until reproductive maturity, but that the survival value for arrangements exceeding this longevity is considerably diminished (Hayflick 2007a).

If aging is driven by thermodynamics, as suggested by Hayflick and further supported here, then any and all factors that contribute towards either resisting or promoting permanent thermodynamically-induced changes in any biocomponent subject to irreversible loss are implicated in longevity determination. This includes factors that directly or indirectly affect the magnitude of the thermodynamic stresses on these biostructures as well as factors that specify redundancy levels, which can offer varying degrees of protection from permanent information loss.

### 8.1 Investigating the Rate of DNA Informational Fidelity Loss in Individuals of a Species

The loss of DNA informational fidelity is inevitable in any individual given sufficient time. If such loss is paramount to aging, then a closer examination of the thermodynamics affecting DNA molecules is warranted and may assist in identifying primary longevity determinants.

Since DNA undergoing replication is significantly more likely to incur a mutation due to the impaired stability of single-stranded DNA (Frederico et al. 1990) and given the imperfect nature of DNA polymerases (Arias-Gonzalez 2012), replicating and non-replicating conditions should be considered independently. Although damage detection and repair systems correct or eliminate many DNA insults, a certain proportion avoids detection. Fig. 6 depicts a systems flow diagram of this arrangement. DNA actively being transcribed is also more susceptible to mutation (Kim and Jinks-Robertson 2012), but it is generally believed that the vast majority of mutations arise from replication events or random DNA damage. For this reason, transcription is not considered as a separate state condition in this model.

**Fig. 6.**
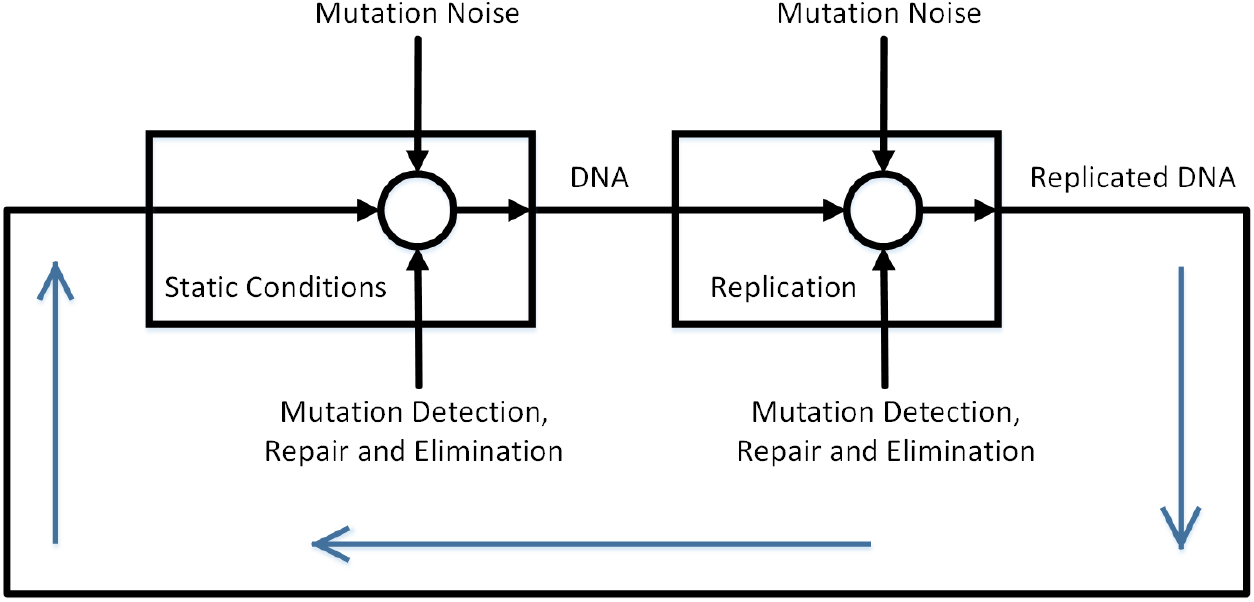
A systems flow diagram of information in a DNA molecular ensemble within a living organism.

Assuming that the time spent in the replicative state is comparatively much less than the time in the static state, a general representation of the average rate of change in DNA information takes the form

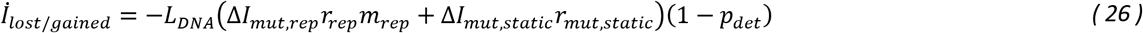

where *L_DNA_* is the length of the DNA molecule in base pairs, *ΔI_mut,rep_* is the amount of mutual information lost or new information gained in the average mutation event during replication and *ΔI_mut,static_* is the same for static (non-replicating) conditions, *r_rep_* is the DNA replication rate, *m_rep_* is the length-specific incidence of mutation during replication, *p_det_* is the probability that a mutation will be detected and eliminated by the cell, and *r_mut,static_* is the length-specific rate at which mutations occur in non-replicating conditions.

### 8.2 Preserving mtDNA Informational Fidelity

Eukaryotic cells (with few exceptions) contain mitochondria, ranging from several hundred to thousands per cell. Each mitochondrion contains at least one copy of mtDNA. Compared to the nuclear genome, the mitochondrial genome is more susceptible to mutation (Larsson 2010) and these mutations are more likely to cause dysfunction. The mitochondrial genome is replicated during mitobiogenesis, which is required for preservation of a healthy pool of mitochondria (i.e. a low degradation state). In any given cell, the mitochondrial pool will be maintained at relatively steady quantities by a combination of mitochondrial fusion, fission, mitophagy, and mitobiogenesis processes; this results in mtDNA replication rates that are very high compared to the rate at which nuclear DNA replicates. Due to the imperfect fidelity of replication with DNA polymerase (Zheng et al. 2006) and the vulnerability of the singlestranded mtDNA replicon (Frederico et al. 1990), each replication event represents a period of time in which the possibility for a mutation is considerably higher than in non-replicating conditions (Kennedy et al. 2013). The microenvironment within mitochondria is also particularly harsh compared to other cellular compartments due to the relatively high concentrations of ROS (Wallace 1999), resulting in larger internal entropy-producing thermodynamic potentials and higher molecular insult rates. Furthermore, as the mitochondrial genome has evolved to be extremely compact, even single-base alterations in mtDNA are likely to cause mitochondrial dysfunction.

The only known human mtDNA polymerase, DNA polymerase *γ*, is highly conserved across species as diverse as *Drosophila melanogaster* and *Saccharomyces cerevisiae* (Chan and Copeland 2009), as are the GTPases implicated in mitochondrial fission and fusion (Ashrafi and Schwarz 2012). Nuclear DNA repair pathways are also highly conserved (Gredilla et al. 2010); mitochondria possess many of the same repair mechanisms and share some of the nuclear DNA repair enzymes. PTEN-induced putative kinase protein 1 (PINK1) and the E3 ubiquitin ligase parkin regulate mitophagy in many metazoans and have homologs across species as diverse as humans and *Drosophila melanogaster* (Cookson 2012). These similarities suggest that the length-specific frequency of a mutation event during mtDNA replication *m_rep_* and the probability that a mutated mtDNA molecule is detected and eliminated or repaired *p_det_* are comparable across a wide range of species.

In addition, since the molecular configuration of DNA is conserved and the potential reactions that can result in molecular modifications are accordingly similar, it follows that the mutual information lost or new information gained in the average mutation-causing event is consistent; i.e. *ΔI_mut,rep_* and *ΔI_mut,static_* should be similar across species. This leaves the mtDNA replication rate *r_rep_* and the static-condition mutation rate *r_mut,static_* as the likely factors from Eq. *(26)* to be primarily responsible for any variation in the rate of mtDNA informational fidelity loss between species.

### 8.3 MtDNA Informational Fidelity Loss in Aged Organisms Primarily Results from Replication

MtDNA mutations increase in an age-dependent manner. High-sensitivity sequencing of human brain tissue from young and old individuals found that most mtDNA point mutations are transition mutations (Kennedy et al. 2011), consistent with replication errors. In addition, 90% of all age-related mutations in mtDNA from human colon are transitions (Greaves et al. 2012). The mtDNA mutation burden in aged *Drosophila melanogaster* is similar to vertebrate levels and also demonstrates a prevalence of transition mutations (Itsara et al. 2014). G:C to T:A transversions, which are typical of oxidative damage, only represented a small percentage of the mutations in these studies.

MtDNA mutation patterns display strand asymmetry consistent with spontaneous cytosine deamination on the lagging strand template during replication (Frederico et al. 1990) in both aged human brain (Kennedy et al. 2011) and aged somatic and germline cells of *Drosophila melanogaster* (Haag-Liautard et al. 2008; Itsara et al. 2014). Mitochondrial mutational spectra produced with purified human DNA polymerase γ accounted for 83% of the mutations found *in vivo* (Zheng et al. 2006). These data strongly suggest that: 1) the vast majority of mutations in mtDNA result from errors during replication; 2) the rate of mtDNA informational fidelity loss varies across species but total losses are similar at species equivalent ages; 3) oxidatively damaged mtDNA is repaired or eliminated with very high efficiency; and 4) oxidatively damaged mtDNA accounts for only a small percentage of mtDNA mutations occurring with age. These postulates are at odds with theories that implicate ROS levels and the resulting direct oxidative damage to DNA as a primary causative factor in aging.

A logical deduction from this is that mtDNA replication rate is higher in shorter-living animals. Unfortunately, the availability of data to support or refute this assertion is limited. Measuring the mtDNA turnover rate *in vivo* has historically proven difficult, although more recent techniques have overcome some of the issues (Collins et al. 2003). Primary cell cultures are required for deriving accurate mtDNA replication rates *in vitro*. Surprisingly, no comparative studies quantitating mtDNA replication rates across a range of species have been published.

### 8.4 Higher Mitochondrial Performance Gives Rise to an Increased Rate of mtDNA Informational Fidelity Loss

As it facilitates mitochondrial renewal, mitobiogenesis is required to preserve the performance of individual mitochondrial components and mitochondria as a whole. Reducing mitobiogenesis will compromise mitochondrial performance since less negative entropy is produced for counteracting degradative internal entropy production (affecting mitochondrial components other than mtDNA), resulting in a shift to a higher degradation state. As mitobiogenesis incorporates mtDNA replication, a decline in mitobiogenesis will reduce mtDNA replication rate *r*_rep_. Thus, although reduced mitobiogenesis may negatively impact mitochondrial performance, it will lead to a lower rate of mtDNA informational fidelity loss per Eq. *(26)*.

On the other hand, with increased mitobiogenesis additional negative entropy is available for offsetting degradative internal entropy production, effectively translating to a lower mitochondrial degradation state and higher performance. However, this will also raise the mtDNA replication rate *k_rep,mtDNA_* and generate increased exposure of mtDNA to the high thermodynamic-stress conditions experienced during replication—resulting in an increase in the rate of mtDNA informational fidelity loss.

Preservation of a mitochondrial degradation state representative of youthful homeostasis requires that the rate of negative entropy production from mitobiogenesis equals or exceeds the rate of degradative internal entropy produced within mitochondria when operating at a youthful degradation state (per Eq. *(7)*). If the mitochondrial performance requirements for a particular cell type are greater in one species, then the rate of mitobiogenesis required to preserve youthful mitochondrial homeostasis within the cells of that species will also be greater, as will the rate of mtDNA informational fidelity loss.

### 8.5 A Closer Look at Mitochondrial Configurations and Membrane Composition

Since mitobiogenesis encompasses mtDNA replication—which accelerates losses in mtDNA informational fidelity—forfeiture of youthful mitochondrial homeostasis is not only inevitable but also temporally related to the rate of mitobiogenesis. How then might this rate differ by species, and why?

Examining cellular metabolic demands provides some clues. Across species, whole-organism basal metabolic rate scales allometrically with body mass: 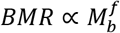 (Kleiber 1932; Peters 1986; Niklas 1994)^8^. Resting oxygen consumption expressed per unit body mass scales proportionally with 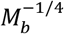. In other words, mass-specific BMR decreases by approximately 16% across species for every doubling of body mass. The inverse correlation between relative oxygen consumption and body mass has been verified in isolated hepatocytes from mammals (Porter and Brand 1995) and birds (Else 2004) as well as in mammalian liver slices (Couture and Hulbert 1995). Porter and Brand (1995) found a 5.5-fold decrease in hepatocyte oxygen consumption for every 12,500-fold increase in body mass.

In general, cells from smaller species have increased oxygen consumption and ATP turnover rates compared to cells from larger organisms. As a result, cells from smaller species place greater energetic demands on their mitochondrial networks. Mitochondrial count correlates with mass-specific changes in tissue metabolic rate (Smith 1956). However, the differences in mitochondrial number per cell cannot fully explain the variation in respiration rate with body mass (Porter and Brand 1995).

Increasing the mitochondrial inner membrane surface area per unit volume of mitochondrial matrix allows for additional transmembrane-localized oxidative phosphorylation enzymatic machinery in the same volume of space (i.e. an increase in *ρ*). The resulting increase in biological power density may help smaller organisms realize their higher ATP requirements. Indeed, a significant negative correlation has been found between mitochondrial inner membrane surface area per unit volume of matrix and body mass (Porter et al. 1996).

Mitochondrial membrane phospholipid composition also differs widely across species, particularly in fatty acid composition (Daum 1985). Smaller mammals have mitochondrial membranes that are more polyunsaturated than larger mammals (Porter et al. 1996). Some light was shed on why membrane fatty acid composition varies allometrically when the molecular activity of transmembrane proteins was examined in different membrane compositions. The cytoplasmic membrane-localized sodium pump (Na^+^·K^+^-ATPase) varies in molecular activity from approximately 8,000 ATP/min in mammals compared to 2,500 ATP/min in ectotherms (all data taken at 37°C) (Else et al. 1996). Cytoplasmic membrane crossover studies demonstrated that the work rate of ectothermic sodium pumps increased significantly when transferred to mammalian membranes, while mammalian sodium pump activity was attenuated in ectothermic membranes (Else and Wu 1999).

The higher sodium-pump work rates in endotherms were hypothesized to be due to influences from surrounding lipids, with polyunsaturated membranes promoting increased molecular activity compared to membranes that are more monounsaturated. Hulbert and Else (1999) proposed a mechanism by which this may occur: The lateral diffusion coefficient of lipids within a membrane bilayer is greater than that of transmembrane proteins by two orders of magnitude (Storch and Kleinfeld 1985). As such, membrane proteins are continuously colliding with membrane lipids. The kinetic energy exchanged during these collisions is believed to be critical in facilitating membrane protein function. The acyl chains of saturated and monounsaturated fatty acids are more flexible than polyunsaturated fatty acids. Therefore, collisions involving lipids containing polyunsaturated fatty acids transfer more energy to membrane proteins and result in higher protein activity than collisions with lipids with only highly saturated fats. Of the fatty acids found in membrane lipids, docosahexanoic acid (DHA or 22:6 n-3) contains the largest number of evenly spaced double bonds but is also particularly susceptible to peroxidation. DHA has been referred to as the "acme" of polyunsaturates and may serve as a membrane "energizer" (Hulbert and Else 1999), effectively increasing *Ẇ_m,max_* (see section 2.8). Sodium pump molecular activity correlates with membrane DHA concentration in both ectotherms (Turner et al. 2005) and endotherms (Turner et al. 2003).

Peroxidation index (PI) is a measure of the susceptibility of membrane lipids to peroxidation and is correlated to fatty acid unsaturation. The PI of mitochondrial phospholipids, predominantly driven by DHA content, negatively correlates with MLSP (Pamplona et al. 1998). Importantly, the same trend line holds for both mammals and birds (Hulbert et al. 2007). In addition, mitochondrial membrane remodeling resulting from various levels of caloric restriction in mice produced changes in PI and MLSP that fit the same trend line (Faulks et al. 2006; Hulbert 2008).

In addition to the negative allometry of metabolic rate, body mass positively correlates with MLSP (Calder 1984; Austad 2005; de Magalhães et al. 2007). The discussed findings suggest that smaller organisms with reduced longevity may utilize membranes with more polyunsaturated membranes—largely dictated by DHA content—in order to increase the rate of work that can be performed by each transmembrane protein molecule *(Ẇ_m,max_)* and to satisfy functional requirements largely specified by recognized allometric relationships that characterize fitness optimization across species. A downside of polyunsaturated fatty acids is their susceptibility to oxidative damage. In other words, polyunsaturated fatty acids are less resistant to molecular alterations resulting from the thermodynamic forces of their environment; the presence of higher levels of polyunsaturated fatty acids will lead to increased rates of degradative internal entropy production *(Ṡ_i_)* within mitochondria—even when mitochondria are functioning well below their maximum work rate (i.e. *w_dc_* is low). This will necessitate a higher rate of mitobiogenesis to maintain homeostasis at a given mitochondrial degradation state.

### 8.6 Identifying Longevity Determinants

It is postulated here that peak biological power density *σ_peak_* largely stipulates membrane composition and other defining characteristics of an organism (e.g. biomolecular renewal rate, metabolism) that serve to largely determine longevity in many species by defining the rate of loss of DNA informational fidelity in somatic cells of the individual. This concept is depicted in Fig. 7. Here the term "peak biological power density" represents the maximum localized volume-specific rate of external work (power per unit volume) achievable within an organism. The cells or tissues where this potentiality exists may vary by species (for example, skeletal muscle in some organisms, neurons in others, etc.). "External work", in the context of peak biological power density, refers to the sum of the biomechanical, biochemical and bioelectrical work that is brought to bear on the immediate environment surrounding the localized region where this work originates. Examples include the mechanical work generated by myocytes, the chemical and electrical work produced by neurons, and the chemical work performed on metabolized products by hepatocytes. External work does not include work associated with housekeeping or "overhead" cellular processes such as biomolecular repair and replacement, maintenance of baseline membrane potentials or mitotic cell turnover.

**Fig. 7.**
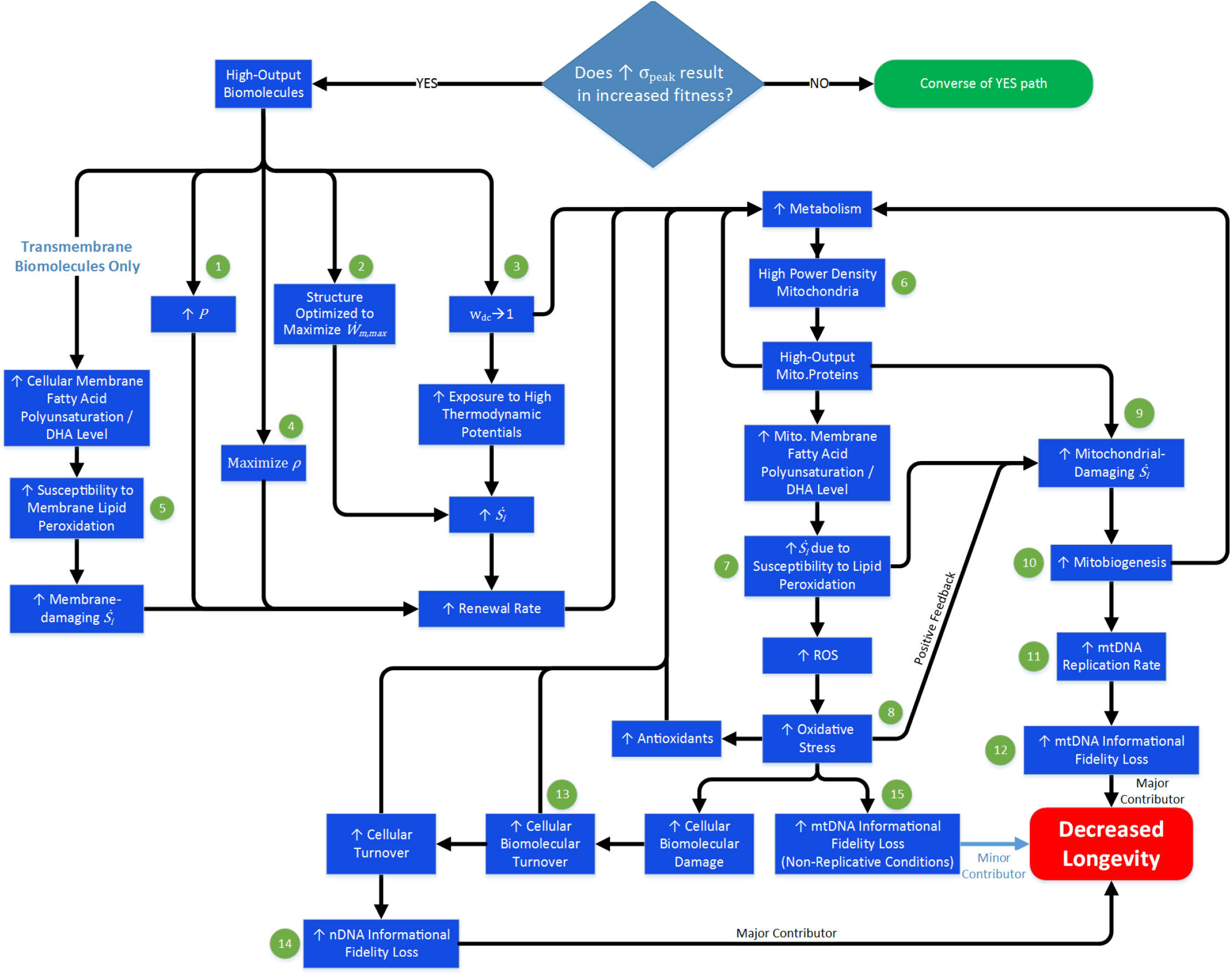
A theoretical means by which an organism's peak biological power density may influence longevity. Bold numbers in section 8.6 text refer to this figure.

To illustrate the sequence of interactions implicated in this model, we will consider an arbitrary organism in which maximum fitness is achieved by a high level of peak biological power density compared to some reference organism (Fig. 7). This implies an increase in the maximum potentiality for external work rate per unit volume that is achievable by some cell, or group of cells, within the organism. Per Eq. *(18)*, in order to realize the highest rate of work possible from a given volume several parameters should be optimized. For one, *P* should be maintained at a high level, **1** (bold numbers in this section refer to Fig. 7). High *P* will increase renewal rate and decrease renewal ROI (Fig. 1D). Secondly, biomolecule structure should be optimized for high peak work rate *(Ẇ_m,max_)*, **2**. This will likely correspond with a reduction in stability (due to lower selective pressure on this parameter), leading to an increase in *Ṡ_i_* and renewal rate. Thirdly, high-output biomolecules should spend more time performing work and less time in a resting state (i.e. *w_dc_ →* 1), **3**. The state of actively performing work involves the transfer of energy and conditions of higher thermodynamic potentials compared to the non-working (resting) state. This will also increase *Ṡ_i_* and renewal rate. Finally, the biomolecule's volumetric density should be as high as possible without negatively effecting the work rate per molecule, **4**. An increased renewal rate will be required to maintain *P* in a larger biomolecular pool. The above factors all contribute towards increased metabolism and ATP requirements. This is consistent with the increased metabolic rates found in smaller animals and the fact that smaller animals renew protein at faster rates compared to larger species (Fig. 3).

Transmembrane proteins exhibit increased activity in membranes containing lipids with high polyunsaturated fatty (DHA) content (Else and Wu 1999; Turner et al. 2003; Turner et al. 2005). Configurations that call for a high level of peak biological power density are likely to use these types of membranes, which are also more susceptible to lipid peroxidation, **5**. This contributes towards an elevated rate of membrane-damaging *Ṡ_i_*, more frequent membrane lipid renewal, and increased biomolecular renewal rate.

The aforementioned increased metabolic requirements generate a need for mitochondrial networks capable of satisfying these higher ATP demands. This is realized with high power density mitochondria, **6**, the characteristics of which are outlined in Fig. 8. High-output mitochondrial proteins optimized for maximal ATP production are expected with these configurations. Similar to their cytoplasmic counterparts, high-output mitochondrial proteins will exhibit increased *Ṡ_i_*. Higher mitochondrial membrane polyunsaturated fatty acid (DHA) content will increase peak ATP output through enhanced transmembrane protein activity but mitochondrial membranes will be more susceptible to peroxidation, **7**. Together with a positive feedback effect from elevated ROS and oxidative stress levels, **8**, this will further increase mitochondrial-damaging *Ṡ_i_*, **9**. A higher rate of offsetting negative entropy production will be required in order to maintain mitochondrial quality and preserve youthful homeostasis. This need can only be realized through an upregulation of mitobiogenesis, **10**, which increases mitochondrial renewal rate but will coincide with a higher mtDNA replication rate, 11, and an increased rate of mtDNA informational fidelity loss, **12**. As mtDNA informational fidelity declines with age, the ability to produce usable energy is compromised and worsens progressively. This generates a downregulation of cellular processes which could largely be responsible for the aging phenotype in many organisms.

Increased oxidative stress in high power-density mitochondrial configurations may elevate thermodynamic potentials in other parts of the cell, **13**. This and the other aforementioned contributors to increased biomolecular damage and renewal rates could be expected to increase the rate of cellular turnover. The rate of nuclear DNA informational fidelity loss will be heightened due to elevated replication rates, increasing the rate at which viable stem cells are depleted, **14**. Increased oxidative stress may also influence the rate of mtDNA informational fidelity loss in non-replicative conditions, **15**. However, due to reasons already discussed, this contribution is probably small compared to the effects from an increased replication rate.

The loss of DNA informational fidelity is unavoidable. Notably, the logic established here describes how the rate of loss of DNA informational fidelity may be a function of an organism's peak biological power density requirements, at least in part. As this rate may be critical in determining the amount of time that passes before youthful homeostasis can no longer be sustained, a potential link is herein established between an organism's peak biological power density and longevity; by this token, peak biological power density could be thought of as a high-level longevity determinant.

Although higher *σ_peak_* implies the ability to perform work at an increased rate per unit volume somewhere within an organism, it does not mean that *Ẇ_m,max_* is achieved at all times or otherwise define *w*_dc_. On the other hand, if this maximum work rate were never approached then there would be no need to possess this ability in the first place. Thus, it is reasonable to expect that *Ẇ_m¡act_* in a species with a higher *σ_peak_* is greater, on average. At a minimum, the cell must have the ability to provide sufficient energetic resources to satisfy *σ_peak_* even if these levels are only attained sporadically. The attributes associated with this last requirement are likely to increase *Ṡ_i_*, renewal rate and thereby DNA informational fidelity loss even in static (non-working) conditions. Thus, the potentiality implied by an increased *σ_peak_* may be sufficient to decrease longevity, even if *Ẇ_m,act_* is lower on average compared to some other species. This could manifest as species with basal metabolisms that are inconsistent with expected longevity trends.

### 8.7 The Naked Mole-rat Paradox - Part II

We can now propose a solution for the second half of the naked mole-rat paradox discussed earlier. Naked mole-rats live in very hypoxic environments and thus must function at extremely low metabolic rates. This necessitates low-output proteins, as high-output proteins have increased metabolic requirements for a number of reasons (Fig. 7). The situation is therefore the reverse of the high peak biological power density scenario previously discussed. Lower metabolism will lead to mitochondria better optimized for stability (Fig. 8). Consistent with this notion, naked mole-rats have one fifth of the amount of DHA in their liver mitochondrial membranes compared to their similarly-sized cousin the house mouse (Mitchell et al. 2007). Decreased susceptibility to lipid peroxidation lowers the rate of damaging internal entropy production, mitobiogenesis, and mtDNA informational fidelity loss. Cellular turnover and the rate of nuclear DNA informational fidelity loss will also decrease. The lower rate of DNA informational fidelity loss increases the amount of time that passes before youthful homeostasis is lost due to transitions to higher degradation states. As a result, the naked mole-rat exhibits exceptional longevity for its size. The increased level of oxidative damage does not limit longevity as it is not the factor forcing a shift from youthful homeostasis but is merely indicative of the high degradation states that coincide with prioritizing renewal ROI for maximizing evolutionary fitness in very hypoxic conditions. I postulate that the exceptional longevity of the naked mole-rat is primarily a byproduct of the aforementioned requirement for extremely low metabolic rate as opposed to direct selective pressure for extreme longevity.

**Fig. 8.**
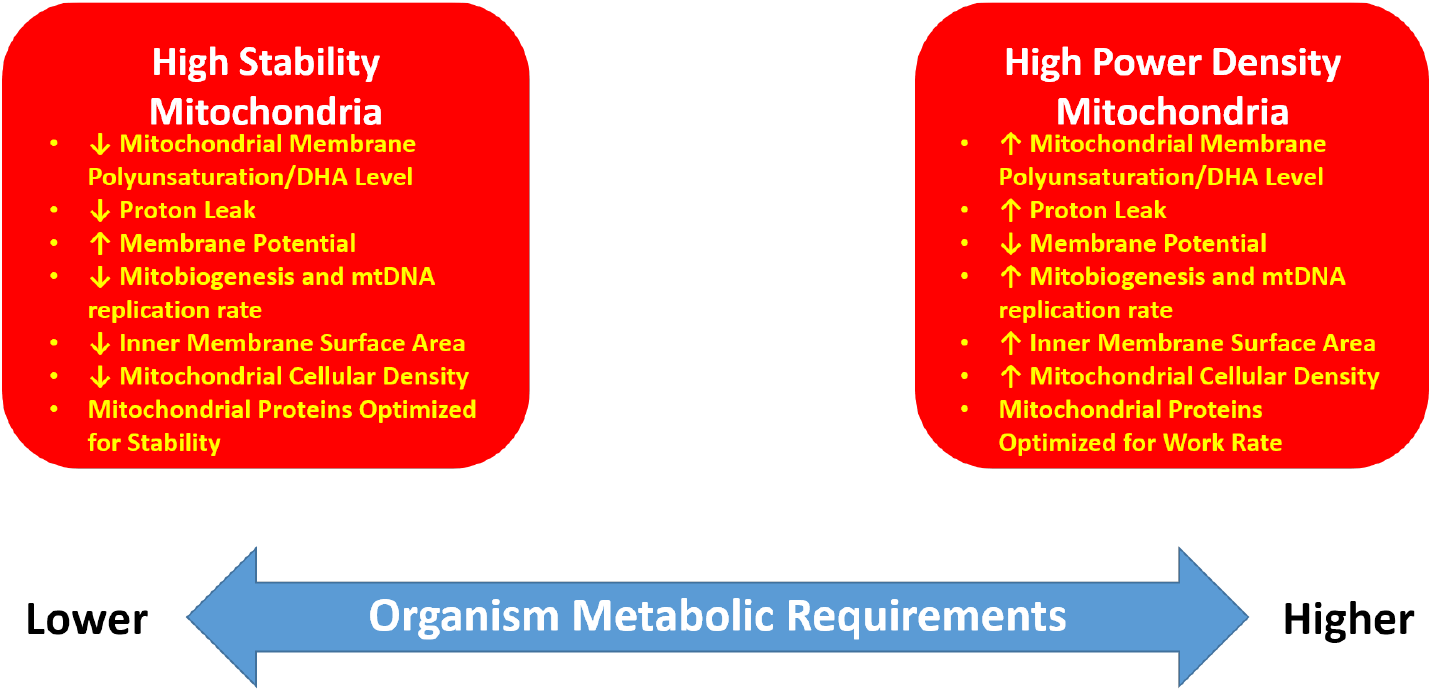
The characteristics of high stability mitochondria compared to mitochondria optimized for peak biological power density. The requirements of the organism dictate where a particular species falls within the range of configurations between these two extremes.

### Allometric Relationships Describe Peak Biological Power Density Trends that Largely Predict Longevity

If peak biological power density is a primary longevity determinant, then how and why does this vary by species? Do variations in peak biological power density align with allometric trends? Some answers to these questions may arise from examining how an organism's mass-specific energetic cost of transport (COT) is driven by certain factors. COT is a measure of the quantity of metabolic energy required to move one unit mass of an organism over one unit distance. In terrestrial animals, COT negatively correlates with body size (Taylor et al. 1982; Strang and Steudel 1990; Reilly et al. 2007). The reasons for the increased locomotor costs in smaller terrestrial organisms have been discussed in detail elsewhere, including Reilly et al. (2007), and Kilbourne and Hoffman (2013). We will briefly examine some of the more significant causes here. Although the mass-specific metabolic energy consumed per stride remains constant across large and small mammals at the same stride frequency, larger animals require fewer strides to cover an equivalent distance; this at least partly explains the reduction in COT with increasing body size (Heglund et al. 1982; Heglund and Taylor 1988; Kram and Taylor 1990). The effect is compounded by the fact that larger mammals have disproportionately longer limbs (positive allometry) (Pontzer 2007).

In general, smaller animals cannot simply decrease their top speeds to offset the increased COT and preserve a low metabolic rate since they must be able to achieve speeds sufficient to evade larger predators. This is demonstrated by the fact that, although top speed does increase with body mass in mammals (Garland 1982), the allometric scaling factor only partially counteracts the increased COT in smaller mammals. In other words, the rate of mass-specific metabolic energy consumed by smaller mammals to achieve their top speed is greater than that of larger mammals.

Posture can also significantly affect COT (Biewener 1989). Smaller terrestrial animals tend to have limbs that are more abducted and flexed during movement (Reilly et al. 2007). Larger animals have more upright postures, which confers a mechanical advantage to anti-gravity muscles. This means that smaller mammals have increased muscular energetic demands for counteracting the flexing moment of the ground reaction force. Larger animals are also able to benefit more from elastic storage since the capacity to store energy in tendons positively correlates with tendon cross-sectional area (Biewener et al. 1981; Bennett et al. 1986; Biewener and Blickhan 1988). Pendular savings can reduce the metabolic cost of locomotion and become increasingly relevant as body size increases in erect animals—but are insignificant in smaller crouched animals (Reilly et al. 2007).

For the above reasons, smaller terrestrial animals have higher peak metabolic rates in their skeletal muscles and supporting organs (heart, lungs, etc.). As skeletal muscle is the major contributor to non-resting metabolism, it should not be surprising that field metabolic rate (FMR) scales with negative allometry (Nagy 2005). This also suggests that peak biological power density is likely to positively correlate with skeletal muscle metabolism.

Surface area scales as a function of body mass per the relation 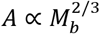. The exponent in this case is less than one, signifying that the mass-specific capacity for heat exchange decreases as body size increases. Since no thermodynamic process is 100% efficient, a portion of the energy utilized for metabolism is converted to heat. The efficiency of the oxidative phosphorylation machinery in mitochondria is highly optimized and not a function of body mass, as indicated by the fact that ATP turnover per unit of consumed oxygen does not change with body mass in mammals (Porter and Brand 1995). Therefore, in the absence of other limiters, maximum sustainable mass-specific metabolic rate will be lower in larger organisms due to their reduced relative capacity to shed metabolic waste heat. In other words, smaller organisms are capable of functioning with higher peak biological power densities. A converse effect of the surface area to mass ratio limits the minimum attainable body size of endothermic amniotes: maintenance of a constant body temperature, below a particular body size for a given set of environmental living conditions, will require an increasing proportion of metabolic energy as body size decreases.

Another factor likely further increases metabolic requirements in smaller animals and limits the minimum attainable body size. West and colleagues argued that metabolic rate scaling is constrained by characteristics of the circulatory system (and other fractal networks) that can be explained by principles from fluid dynamics (West et al. 1997; West and Brown 2005). As vessels become smaller, viscosity leads to greater energy dissipation due to increased damping. In addition, at some critical minimum size vessels are no longer able to benefit from impedance matching, which can greatly reduce the energy lost due to reflections in larger vessel branch points. These energy-consuming effects play an ever-increasing role as body size decreases and narrow vessels predominate.

Although the allometric relationships between BMR/FMR and *M_b_*, and longevity and *M_b_*, at the species level are well established, these describe only general trends. Clearly, the described allometric relationships are not imposing a strict, specific value on peak biological power density, metabolic rate, or longevity for a species or individual. Rather, they describe median values, which suggests that these parameters must evolve within upper and lower bounds that are a function of body size, and that the optimal compromise between peak biological power density, longevity and body size for a given species must also fit within these general constraints—but will also depend on a number of other factors. Thus, deviations from the trends are expected. For example, a species living in an environment with low predatory pressure may receive a fitness benefit from sacrificing peak athletic performance for increased longevity. Suppose that in this case, it is not necessary for the organism to function anywhere near the metabolic limit dictated by its capacity for heat exchange in order to ensure high rates of survival and fecundity over its lifespan; it receives more fitness benefit from maintaining a reasonable level of fecundity over an increased lifespan than from a marginal decrease in predation over a shorter lifespan. Another example of an expected significant deviation from the median are situations where organisms utilize a specific behavioral tactic or enhanced cognitive capabilities to increase their survival odds in lieu of maximizing peak biological power density. Humans are the ultimate embodiment of such a strategy.

These generalized allometric relationships do not apply to individuals within a species. For example, larger individuals in many species, such as dogs (Speakman et al. 2003) typically have shorter lives than smaller individuals. This may be in part because longevity determination has evolved, and is genetically engrained, at the species level. In other words, the genetic elements that specify peak biological power density, membrane composition, biomolecular turnover rates, stem cell reserve levels, and other factors that contribute towards resisting (or promoting) permanent thermodynamically induced changes in biocomponents subject to irreversible losses are mostly preset within the genome of a species and do not vary significantly as a function of body size. It is not surprising that significant deviations from the median body size would result in a compromised individual—and that this would include decreased longevity.

## 9 Longevity Optimization

Once mitochondrial dysfunction has progressed to the point that resource deficits prevent the funding of all cellular processes at youthful levels and/or genetic redundancies are no longer able to compensate for losses in nuclear DNA fidelity, an aged phenotype will begin to take shape. It is reasonable to expect that the optimal allocation of resources for preserving maximal survival and fecundity in an aged individual would be different from the configuration used in young adulthood when adequate resources are available to fund all cellular processes. At all life stages, those factors most critical to immediate survival are of highest priority to the individual. Therefore, a genotype optimized for an aging individual would increasingly deprioritize less vital processes and biocomponents as useable energetic resources become scarcer so that biocomponents that are more critical to immediate survival are retained at degradation states sufficient to sustain life and survival potential/fecundity are maximized. As aging continues to progress in the individual, eventually a state is reached where even vital factors cannot be adequately maintained; at this point the individual's overall condition becomes unconducive to continued life.

Could such an anti-aging strategy exist in multicellular organisms? A large number of genetic elements regulating pathways apparently related to longevity have been identified (ENCODE Project Consortium et al. 2007). Some scientists believe that these pathways are directly responsible for causing aging and that they modulate the rate of aging within a species (Kirkwood 2005; Vijg and Campisi 2008; Austad 2009; Holliday 2010). A proposed complementary hypothesis is that longer-living species have evolved to contain superior mechanisms and/or biomolecules for retarding senescence; some scientists believe that incorporation of these changes into shorter-living organisms could lead to delayed senescence in these other organisms as well.

I submit here an alternative hypothesis proposing that a major function of the putative aging pathways is to optimize the inevitable process of aging such that individual longevity and fecundity are maximized. Contained within these pathways, genetic elements that I term "longevity optimizers" work together to elicit a balanced response to the unavoidable progression towards increasing levels of irreversible biomolecular fidelity loss.

To my knowledge, this concept has not been formally proposed previously. There are several likely reasons for this. Firstly, scientists do not generally acknowledge that aging is inevitable, regardless of genotype. Many popular aging theories (e.g. antagonistic pleiotropy, mutation accumulation, and disposable soma) utilize evolutionary concepts to explain the existence of aging and do not consider fundamental physical law as relevant to the issue. These theories claim that aging is not unavoidable but rather that it exists because it projects beneficial effects on species fitness in other ways (antagonistic pleiotropy, disposable soma) or that there is insufficient evolutionary pressure to eradicate aging (mutation accumulation). If fundamental physical law does not mandate biological aging, then there is no need for mechanisms or strategies to resist or optimize it.

Here, I have provided rationale and evidence for why biological aging is an inevitable consequence of fundamental physical law that evolution cannot overcome. If this is the case, then it is reasonable to propose the existence of evolved mechanisms to resist and optimize an organism's susceptibility to these effects in order to maximize species fitness.

A counterargument is that aging optimizations are unlikely to evolve because selective pressures begin to decrease for ages beyond reproductive maturity. However, as the potential for loss of DNA informational fidelity begins at conception—not at reproductive maturity—this phenomenon must be suitably combatted at all life stages. In order to maximize fitness, organisms require strategies for preventing the loss of DNA informational fidelity from reaching detrimental levels and to compensate for the DNA informational fidelity loss that has occurred at all stages of life.

### 9.1 Selective Pressures Favor Genotypes that Attenuate Increases in Mortality and Losses in Fecundity Occurring After Reproductive Maturity

In the absence of compensatory mechanisms, somatic mutations and other forms of irreversible degradation to an essential biocomponent (biomolecule, cell, tissue, structure, etc.) will nearly always have neutral or negative effects on mortality rate and fecundity. Therefore, the integrative effect of the systemic degradation occurring with age must eventually result in tangible negative repercussions for the individual.

In any aging individual organism, biocomponents susceptible to irreversible fidelity loss will be the first to transition from their youthful homeostatic states; therefore, aging of the individual will initiate within these structures. Multicellular organisms with short lifespans, such as many insects, often contain critical structures that grow once and are not maintained. One example is the wings of a honeybee. A honeybee's wings are primarily constructed of dead cells that cannot be replenished. Thus, any damage sustained to the wings will remain throughout the honeybee's life. In contrast, although (in general) mammals lack the ability to repair damage resulting from major traumatic injury to critical structures, they are able to counteract everyday "wear and tear" for considerable lengths of time by way of renewal and repair at the cellular and tissue level.

In the case of organisms with short lifespans and susceptible critical structures, irreversible fidelity loss in such structures may be highly relevant to the initiation and progression of the aging phenotype of these organisms. On the other hand, most organisms with at least moderate longevity will have very few biocomponents that lack fundamental repair capacity. For this reason, in all but the shortest-living multicellular organisms, the susceptible biocomponent most likely to drive the aging process is DNA. The continual loss of DNA informational fidelity must eventually force shifts in the degradation state of other biocomponents.

The magnitude of any deleterious impact on individual instantaneous mortality rate and fecundity resulting from an increase in the degradation state of a biocomponent will depend on the function of the biocomponent and the extent of the shift in degradation state. One such strategy for maximizing survival rate and fecundity in these conditions is to minimize the degradation state, or failure likelihood, specifically of those biocomponents most critical to these parameters. We will examine whether such a strategy would be evolutionarily favored.

Hamilton (1966) exploited the Euler-Lotka equation (Euler 1767; Lotka and Sharpe 1911; Fisher 1930) to derive a measure of fitness *r* from age-specific survival and fecundity rates.

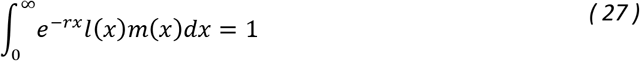

Here *l*(x) represents survival up to age *x* and *m*(*x*) is fecundity at age *x*. Using a similar framework, Fisher (1930) introduced the concept of age-specific reproductive value *v*(*x*) with the following relation

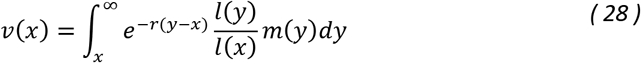

Fisher (1930) described reproductive value as a measure of the contribution of individuals of age x to the future ancestry of a population and stated that (p.27) "the direct action of natural selection must be proportional to this contribution". In other words, genotypes that maximize reproductive value for a given age are evolutionarily favored over those that produce a lower *v*(*x*). Fisher also discussed why reproductive value typically increases before reaching an apex and then declines with age.

Assume that in some hypothetical organism, *m*(*x*) peaks near reproductive maturity before declining, and that mortality increases from this same point forward, accelerating the rate at which *l*(*x*) decreases with age. This age-related decline would be typical of the inevitable and irreversible fidelity losses described above. The red curve in Fig. 9A depicts a typical plot of reproductive value as a function of age for this scenario (calculated using Eqs. *(27)* and *(28)*). This is representative of an organism lacking genetic elements for optimizing fecundity and survival in response to irreversible losses in fidelity. If the losses in *l*(*x*) and *m*(*x*) occurring after reproductive maturity are attenuated, peak reproductive value will increase and occur at a later age, and reproductive value will be maintained longer (Fig. 9A, blue curve). Due to the positive contribution to reproductive value, genes/genotypes that attenuate the described pattern of losses in *l*(*x*) and/or *m*(*x*) will be evolutionarily favored, provided they do not negatively influence early reproductive value.

**Fig. 9.**
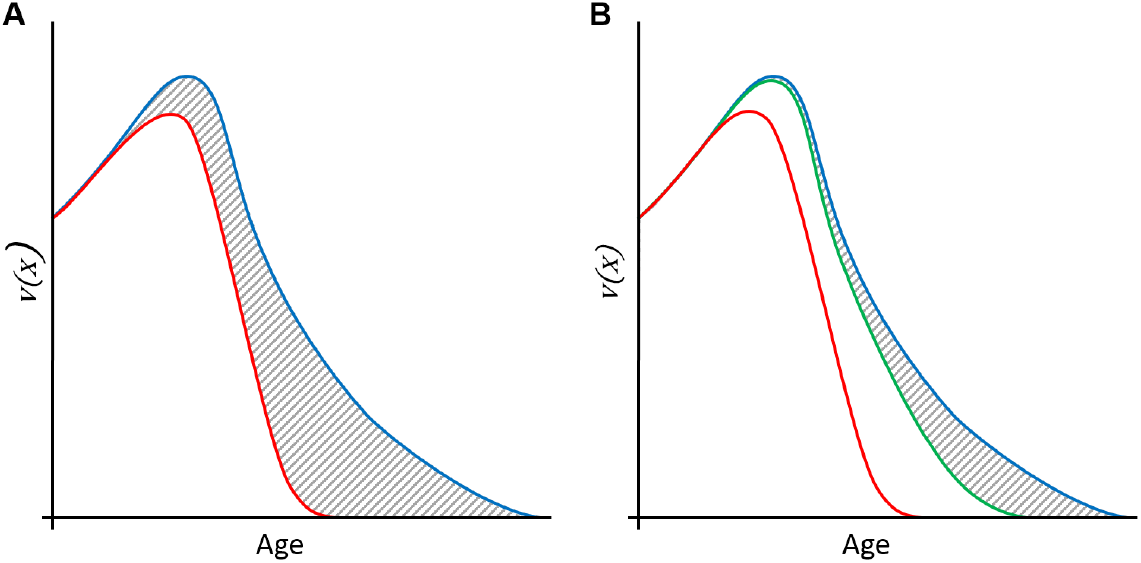
Two scenarios of reproductive value curves for a hypothetical organism. (a) Optimization (blue curve) of the organismal response to irreversible losses in fidelity (as detailed in text) will provide a fitness advantage compared to a genotype lacking longevity optimizers (red curve), due to the increase in reproductive value depicted by the grey shaded region. (b) Variations on optimization. Three genotypes are illustrated: no longevity optimization (red curve), ideal longevity optimization (blue), and partial longevity optimization (green). All curves were modeled according to parameters described in the text and using Eqs. (27) *and* (28). *Curves depict calculated trends*.

### 9.2 Deterioration Management Strategies

It is illogical for an organism to have evolved such that fecundity or mortality is negatively affected (at ages where selective pressure is still above some minimal threshold) due to the disproportionate deterioration, or increased likelihood of failure, of one or a small number of vital biocomponents. Selection must favor genotypes that avoid susceptibility to the catastrophic failure of a small number of weak links during aging.

I propose that cellular mechanisms and pathways have evolved to function in a progressive and dynamic manner to manage the irreversible, and inevitable, losses in fidelity afflicting an aging individual. Priority is placed on biocomponents most susceptible to degradation effects, and most critical to survival and fecundity. To illustrate this concept, I will describe two strategies: "managed deterioration" and "unmanaged deterioration".

In unmanaged deterioration, the degradation of biocomponents occurs at a rate proportional to the biocomponent's susceptibility to irreversible fidelity loss, or the direct and indirect effects of degradation present in other biocomponents (Fig. 10, left). Regardless of their importance to instantaneous mortality rate or fecundity, the most susceptible components reach failure levels first—leading to premature reductions in survival probability and fecundity—while other biocomponents could remain at relatively high performance levels (i.e. low degradation states).

**Fig. 10.**
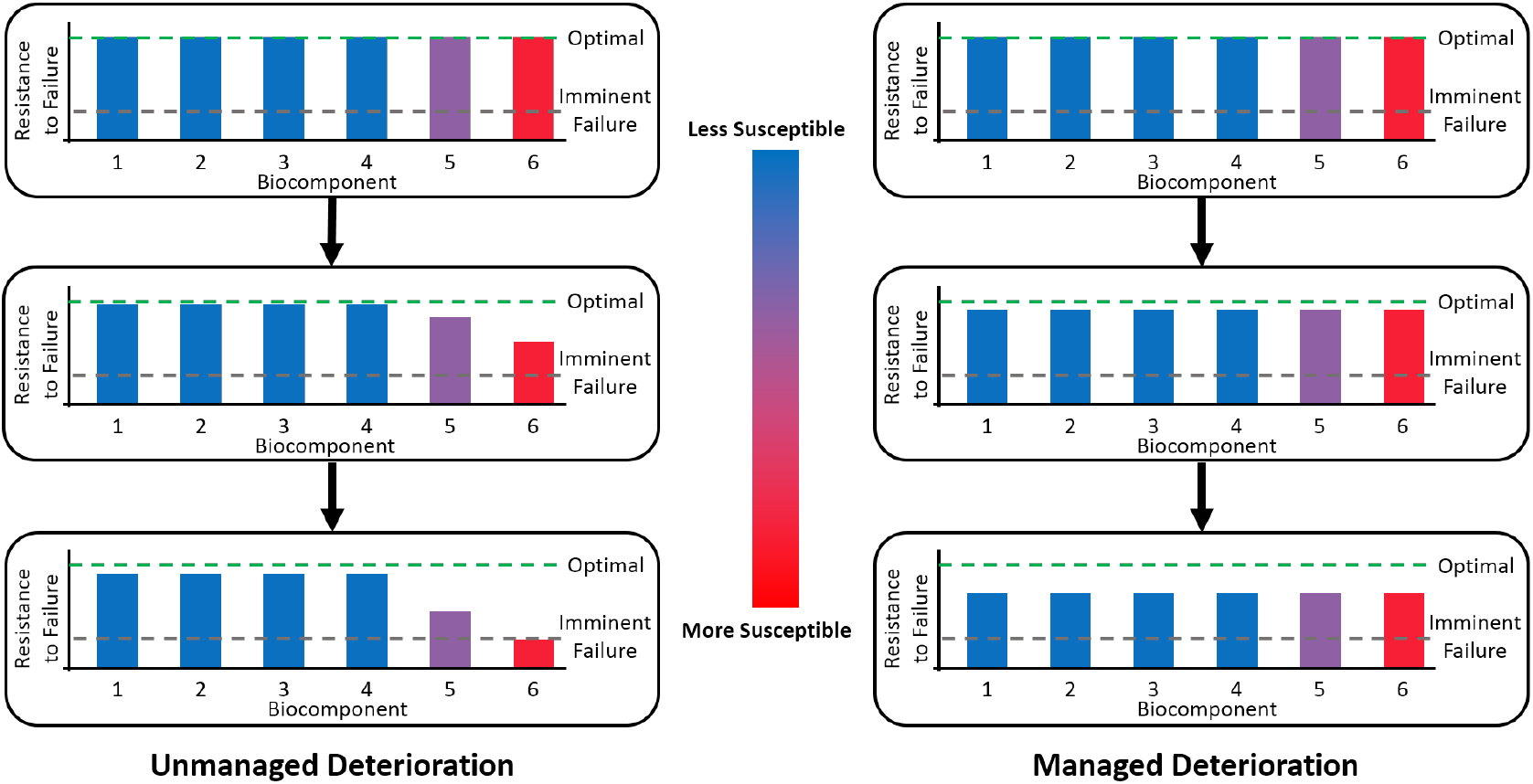
Demonstration of unmanaged (left) and managed (right) deterioration strategies. A group of arbitrary biocomponents, equally vital to an organism's fecundity/survival, are depicted. Three ages are considered: Top—Young Adult, Center—Middle-Age, Bottom—Elderly. 'Red' indicates that a biocomponent is very susceptible to irreversible fidelity loss (which could be due to direct and/or indirect effects), while 'blue' signifies that a biocomponent has very little or no susceptibility to irreversible fidelity loss. The vertical axis depicts how resistant a biocomponent is to failing, given its current degradation state. In unmanaged deterioration, biocomponents will approach imminent failure at a rate proportional to their susceptibility to the effects of irreversible fidelity loss. Biocomponents most susceptible to irreversible fidelity loss will reach failure levels first and the organism will die prematurely. With managed deterioration, longevity optimization genes produce adjustments in the aging individual which partially offset decreases in resistance to failure of the most vital and susceptible biocomponents (details in main text). By not allowing any one vital factor to reach the imminent failure state at an earlier age than others, managed deterioration strategies may enhance longevity and increase fecundity.

In managed deterioration, longevity optimization genes modulate the deterioration rate of biocomponents so that those of similar importance degrade at comparable rates and/or reach their failure threshold at equivalent ages. Critical biocomponents are prioritized. No singular biocomponent is permitted to reach a degradation state that unduly compromises fecundity or survival probability—effectively increasing longevity and overall fitness (Fig. 10, right). This could be accomplished by several means, including:

1. Reallocating resources, at the cellular level and higher, as usable energetic resource availability declines to prioritize biocomponents most important to preserving reproductive value.
2. Adjusting microenvironmental conditions to decrease thermodynamic potentials on more vital biocomponents and thereby lowering the rate of damage-inflicting internal entropy production.
3. Reducing biocomponent turnover rates to delay the clonal expansion of irreversibly compromised biocomponents. (Effectively reduces the rate of fidelity loss.)
4. Altering physiology so that stresses on biocomponents that are more vital are reduced to help maintain these biocomponents within their functional limits.

One possible example of items (2) and (3) is the way by which controlled decreases in mitochondrial fusion and fission may attenuate mtDNA informational fidelity loss during aging. Mitochondrial fusion/fission rates are integral to mitophagy and mitobiogenesis (Twig et al. 2008; Youle and Narendra 2011) and are therefore critical for preserving the performance of mitochondrial components. Reducing fusion/fission compromises mitochondrial quality by increasing the load of ROS products and otherwise-damaged mitochondrial components. Mitochondria in aged organisms produce less usable energy (Yaniv et al. 2013), likely due to the combined effects of reduced mitochondrial fusion/fission and mtDNA informational fidelity loss that occurs with age. Figge et al. (2012) demonstrated that decelerating mitochondrial dynamics actually helps to preserve mitochondrial performance. MtDNA informational fidelity losses are attenuated by reducing the exposure of mtDNA molecules to the high thermodynamic stress conditions encountered during replication and delaying the spread of "parasitic" mutated mtDNA molecules. This is evidently more critical to preserving mitochondrial performance than the tradeoff of increased degradation state (reduced performance) in other mitochondrial biocomponents. In the context of the current discussion, the genes responsible for realizing this strategy are longevity optimizers.

Skeletal muscle mass decreases substantially with advanced age (Grounds 1998). Scientists generally regard this loss as a purely undesirable physical manifestation of aging, and view a safe and effective therapeutic intervention for reducing age-related skeletal muscle mass loss as being desirable and beneficial for the health of the elderly. For example, a substantial body of evidence suggests that the loss of function in satellite cells is a proximal cause of age-related muscle mass loss (Carlson and Conboy 2007; Sousa-Victor et al. 2014) and interventions have been proposed for "correcting" this deficiency (Carlson and Conboy 2007; García-Prat et al. 2013; Sousa-Victor et al. 2014; Sousa-Victor et al. 2015; Dumont et al. 2015).

I offer an alternative hypothesis for explaining this and perhaps other age-linked traits. Given that cardiac output declines significantly with age (Brandfonbrener et al. 1955) and is rooted in functional deficits at the cardiomyocyte level (Guo and Ren 2006), a reduction of skeletal muscle mass will lower the stresses on an age-compromised heart by reducing the volume of blood in the body and decreasing the contractile forces required to circulate the blood. This raises the possibility that a decrease of skeletal muscle mass with age is a beneficial, evolved response—or at least a tolerated condition—which reduces cardiovascular stress and lowers the mortality risk of cardiac events. This is one example of how age-dependent physiological alterations could decrease the likelihood of failure of more critical biocomponents in light of inevitable losses in fidelity, as proposed in item (4) from the above list, and serve to extend longevity. To be clear, this hypothesis is not intended to explain extreme muscle wasting outside of normal age-related trends, which is undoubtedly a genuine pathological condition. In addition, there are certainly a number of undesirable aspects of age-related skeletal muscle dysfunction. The concept put forth here is that age-dependent physiological alterations, even those that at first glance appear purely detrimental, may actually serve a purpose in establishing a balanced configuration in the face of inevitable, and progressively increasing, fidelity losses.

It is prohibitively difficult to prove that the altered age-dependent expression of one gene represents an evolutionarily established tradeoff with some other gene(s) that extends longevity, as suggested by item (1). However, beyond the evolutionary argument for their existence, there is other evidence suggesting that mechanisms of this type might exist—specifically, features of the proteomic, gene expression, and epigenetic signatures in aging individuals.

Gene expression signatures display a characteristic age-associated pattern of changes in specific genes in mice, rats and humans which is consistent across multiple tissue types (de Magalhães et al. 2009), demonstrating conservation of aging signatures across species.

For example, lysosomal genes are overexpressed with age. It is plausible that the decreased protein turnover exhibited in an aged individual could lead to a greater load of proteins that have degraded to the extent that they cannot be processed by proteasomes and therefore they must undergo lysosomal degradation. Although the energetic resources dedicated to increased lysosomal expression could have been allocated to lessening the severity of the general reduction in protein turnover, it may be that age-dependent lysosomal overexpression optimizes the overall protein degradation state based on energetic resource availability and that this balance represents the best compromise for maximizing reproductive value.

DNA methylation patterns also change in line with chronological age in humans (Christensen et al. 2009; Boks et al. 2009; Rakyan et al. 2010; Bocklandt et al. 2011; Bell et al. 2012; Heyn et al. 2012; Gentilini et al. 2012; Garagnani et al. 2012; Horvath 2013; Hannum et al. 2013; McClay et al. 2013; Florath et al. 2013; Christiansen et al. 2015). Predictors can reliably estimate the age of human cells from any human tissue type based on epigenomic DNA methylation profiles (Horvath 2013; Hannum et al. 2013). This supports the notion that age-related epigenetic signatures do not simply represent accumulated regulatory dysfunction, but that at least some component of this signature represents a progressive and dynamic response to aging.

### 9.3 Longevity Optimization Strategies from Early Adulthood May Serve as Templates for Those Used in Later Life

The concept advanced here proposes that selective pressures have led to the evolution of genetic optimizations that attenuate the rate of loss of DNA informational fidelity (and irreversible fidelity losses in other susceptible biocomponents) and the detrimental effects of these losses in aging individuals. There can be little doubt that in the face of inevitable, irreversible and progressive fidelity losses, a diverse array of intermediate configurations would be required to realize optimal aging during all stages of life. As compromised biomolecules reach non-trivial levels even during early adulthood (Ben-Zvi et al. 2009; Greaves et al. 2014), it is reasonable to propose that longevity optimizers have evolved to incorporate complex modulatory strategies to help establish a more balanced overall phenotype for each and every stage of life.

This suggests that evolved, early adult-life longevity optimization pathways could serve as a substantive basis for a late-life longevity optimization strategy. It may be largely through the extrapolation of these early adult-life mechanisms that the maximal lifespan of an organism is able to extend well beyond the age of peak reproductive value, particularly in species such as humans where older individuals have the benefit of protected environments. The use of pre-existing genes and pathways as a basis for later-life optimizations may also explain how genetic elements could evolve to a highly optimized state for advanced ages within a relatively short period of time, even though selective pressure decreases with age (Medawar 1952; Williams 1957; Hamilton 1966). If genes and pathways for early adult-life longevity optimization were already present within an organism's genome, the extension of these strategies for late-life longevity benefits would require considerably less selective pressure and could be realized within a fewer number of generations. Yet, since selective pressure declines with age, it would still likely take an extremely long time for late-life longevity optimizations to evolve in order to fully maximize longevity extension potential. For this reason, longevity optimization in all organisms is almost certainly not fully realized.

Incorporating longevity optimization genes capable of maximally attenuating losses in fecundity, and increases in mortality, preserves late-life reproductive value (Fig. 9B, blue curve) compared to a genotype that lacks longevity optimization (red curve). Now suppose that longevity optimization is close to ideal for ages near peak reproductive value but becomes progressively less so with increasing age and decreasing selective pressure. The reproductive value curve for this scenario (Fig. 9B, green curve) is between the two described extremes. This last curve may be representative of the evolved state of the typical metazoan. The gray shaded region represents the "intervention potential"—the maximal gains in reproductive value attainable by further genetic longevity optimizations or through artificial manipulation of individuals (i.e. drugs and therapies, excluding therapies that restore fidelity in biocomponents subject to irreversible loss). Although beyond the scope of the current discussion, by examining statistics of proportionate mortality by pathological condition and other population data, it may be possible to calculate the ideal longevity optimization curve for a particular organism (Fig. 9, blue curve).

### 9.4 Entropy-Driven Managed Deterioration in Further Detail

It is theorized here that metazoans have evolved to make compensatory adjustments as individuals age so as to minimize the deleterious effects of thermodynamic phenomena on reproductive value—resulting in survival, for the moment, but nonetheless unable to avoid an ever-increasing negative phenotype. These longevity optimizers may protect the biocomponents of an organism at all levels (biomolecules, cells, tissues, and organs) that are most critical to immediate survival and fecundity by sacrificing other aspects of health, leading to a diverse "spread the misery" phenotype. In essence, the diversity of the biocomponents affected during aging and the relatively high degree of conservation of the aging phenotype across taxa may largely be manifestations of these compromises. A more detailed depiction of this model links further aspects of the aging process (Fig. 11).

**Fig. 11.**
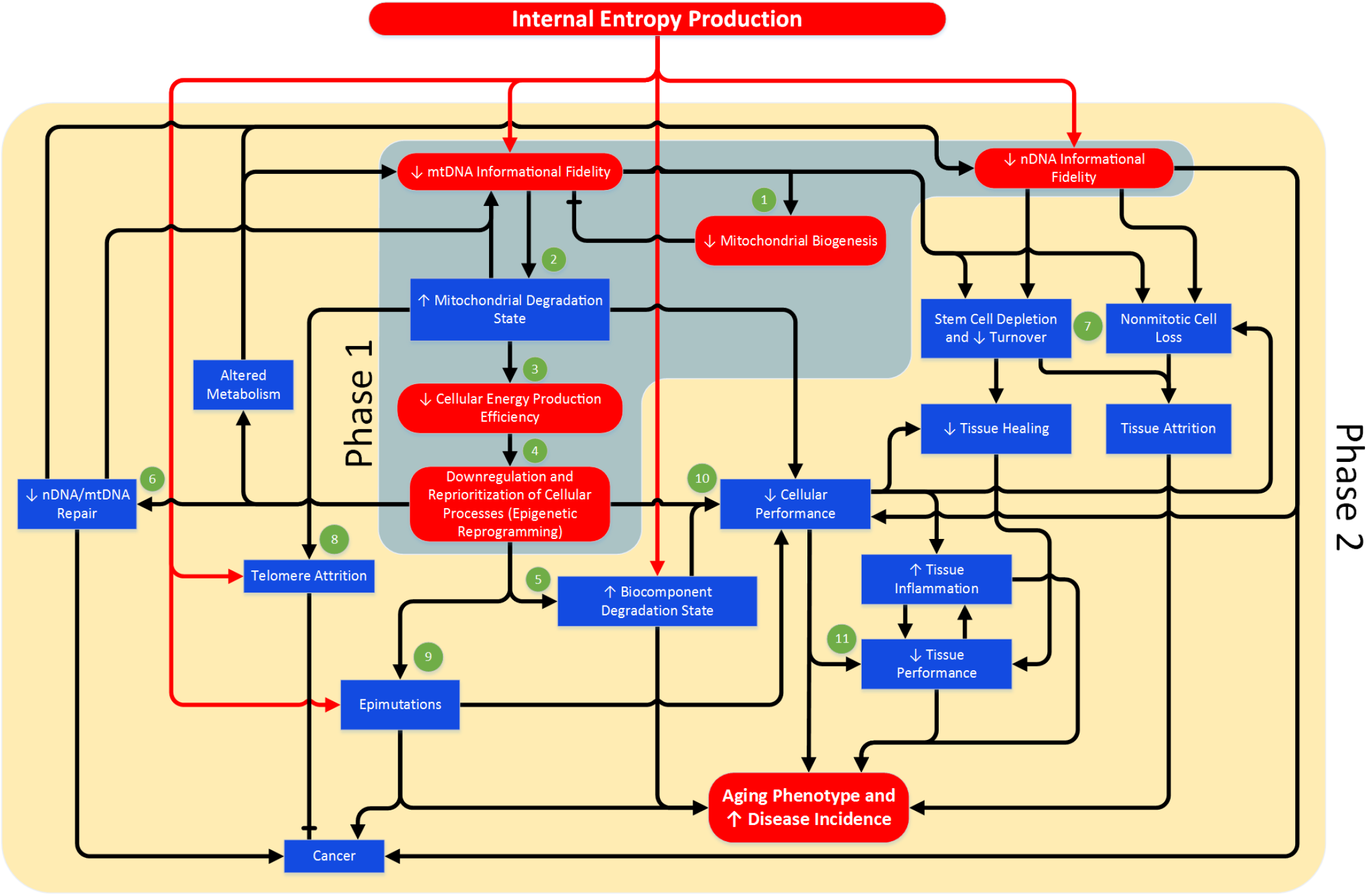
A more detailed look at the higher-level interactions implicated in this theory during the progression of the aging phenotype. The red lines highlight where the degradative effects of internal entropy production are directly exerted. MtDNA informational fidelity loss drives a deceleration in mitochondrial fusion/fission, 1—mitigating, but unable to prevent, further losses. Combined with reduced fusion/fission rates, losses in mtDNA informational fidelity lead to elevated levels of mitochondrial ROS products and compromised mitochondrial components (i.e. increased degradation state), 2, which reduces peak cellular energy (ATP) output (Yaniv et al., 2013), 3. The effects of this are partially tempered by an evolved response that includes changes in resource allocation as well as physiological alterations, 4, and is largely signified by the epigenetic state of a cell. These age-dependent epigenetic signatures should not be confused with epimutations, where distribution is mostly random (Heyn et al., 2012). Mandatory energy conservation reduces the cell's ability to preserve youthful biocomponent degradation states, 5. Nuclear and mitochondrial DNA fidelity is gradually further compromised as renewal processes are downregulated, 6. Increasing losses in nuclear DNA informational fidelity with age (Podolskiy et al., 2016) contribute to increased cell-to-cell stochasticity in gene expression (Bahar et al., 2006) and clonal mosaicism (Lodato et al., 2015), causing average cellular performance to decrease. The loss of DNA informational fidelity also decreases stem cell viability and consumes stem cell reserves, in addition to generating losses in the number and viability of nonmitotic somatic cells, 7. Dysfunctional telomeres can activate the DNA damage response pathway, engaging tumor protein p53 and leading to promotion of apoptosis or replicative senescence (Deng et al., 2008). Telomere attrition is upregulated in aged cells (Passos et al., 2007). This is an evolved mechanism, distinct from the length reduction that occurs during replication, believed to partially offset the increased likelihood of developing cancerous mutations in age-compromised cells (Campisi, 2005), 8. This adaptive response involves the preferential degradation of telomeric DNA in conditions of increased mitochondrial superoxide production (Passos et al., 2007; Petersen et al., 1998; Zglinicki, 2002), as occurs with aging. Epigenome maintenance is downregulated in aged mammals (Cencioni et al., 2013), resulting in an increased number of unrepaired spontaneous epigenome mutations (Chambers et al., 2007), 9. This, combined with escalating DNA informational fidelity losses and the downregulation of DNA damage repair mechanisms (Beerman et al., 2014; Zhang et al., 2010), contributes to an ever-increasing risk of developing cancer (Hansen et al., 2011), as seen with advancing age (American Cancer Society, 2013). Inevitably, the result of cellular component-level degradation is compromised cellular performance, 10, and a concomitant loss in the performance of macro structures: tissues, 11, organs and, ultimately, reduced viability of the organism itself.

## 10 Connecting the Dots

### 10.1 Differentiating between Longevity Determinants and Longevity Optimizers

The model described here proposes that two groups of factors contribute to the intrinsic longevity of a species: 1) longevity determinants and 2) longevity optimizers. It is important to differentiate between these distinct, but occasionally overlapping, groups. Longevity determinants are defined as factors that directly or indirectly influence the basal rate of loss of fidelity in any biocomponent (biomolecule, organ, tissue, etc.) susceptible to irreversible fidelity loss (such as DNA). The genetic arrangements that ultimately determine an organism's basal longevity are driven by fundamental physical law and evolutionary factors, and are further contingent upon the exact environment and environmental interaction factors in which the species exists (Fig. 12). Any genetic element specifying a phenotypic characteristic that influences the basal rate of aging is a longevity determinant, as are the phenotypic characteristics themselves. Macro-level characteristics that may be longevity determinants include peak biological power density, physical size, athletic ability, and metabolic rate. At the micro-level, longevity determinants may include stem cell niche size, membrane composition, biomolecular degradation state, biomolecular durability, and the degree of intramolecular genetic redundancy. Environmental determinants of species basal longevity include temperature/climate, resource availability (food, oxygen, etc.), predation pressure and other factors that mandate tradeoffs between fecundity/mortality and longevity. Survival strategies, and behavior in general, can also influence basal longevity by providing competitive advantages that result in reduced negative repercussions associated with characteristics that serve a role in longevity determination. The subdivision of longevity determinants into genotypic, phenotypic, and environmental elements allows for a clearer depiction of the interplay between the drivers and these different factors.

**Fig. 12.**
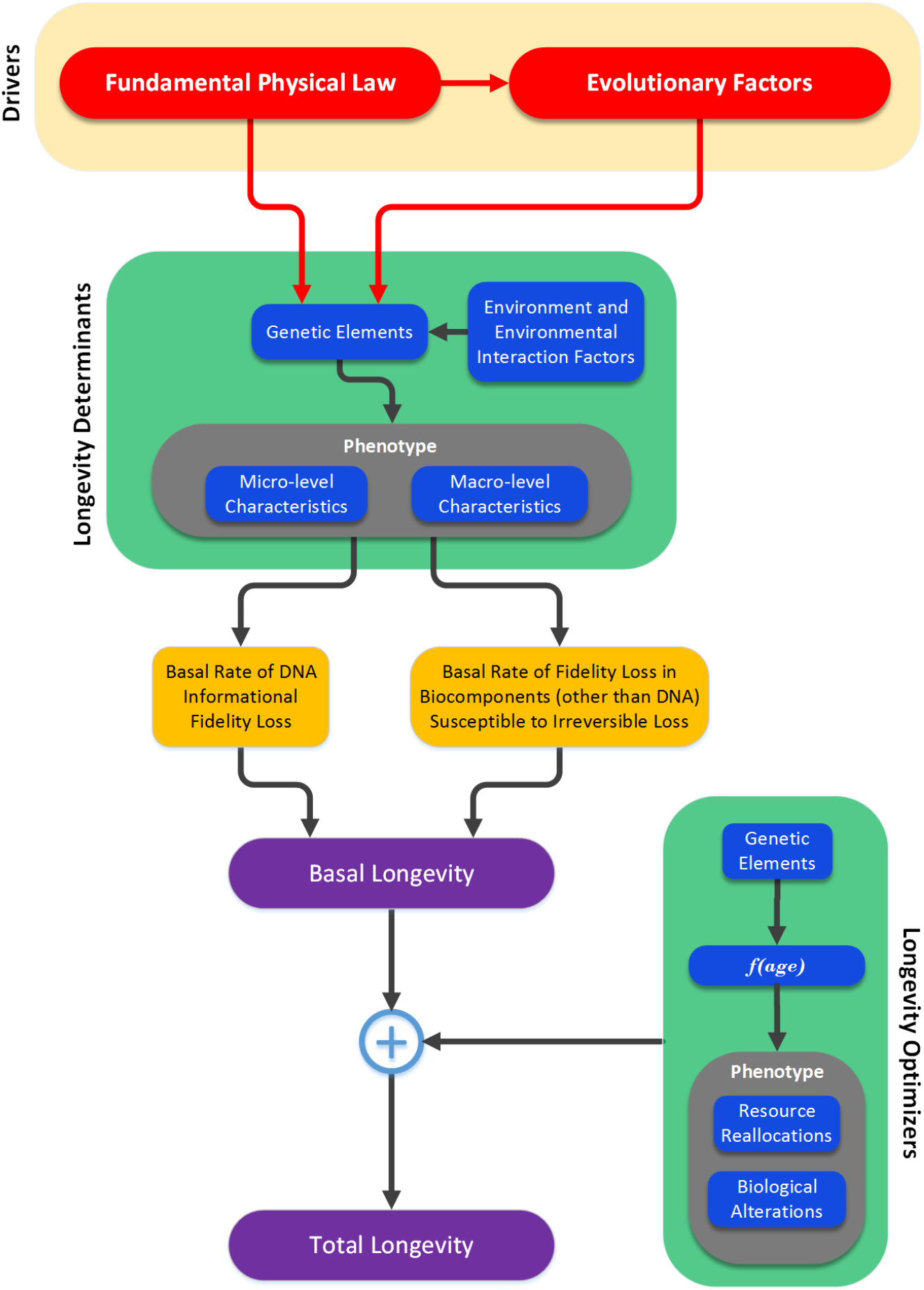
The proposed relationship between longevity determinants, the root causes of aging (explained by fundamental physical law and evolutionary theory), longevity optimizers, basal longevity and total longevity of a multicellular species.

In contrast to a longevity determinant, a longevity optimizer is any genetic element that increases longevity by contributing towards an effect that generally becomes progressively more dominant with age and that delays the severity, or rate of progression, of the aged phenotype. Longevity optimizers reallocate resources and alter physiology so that the overall state of the organism maximizes instantaneous survival rate and fecundity at all ages. To summarize this model, longevity determinants define an organism's basal longevity while longevity optimizers seek to further maximize longevity through dynamic adjustments during the aging process that ultimately serve to balance the aging phenotype.

### 10.2 The Number of Genetic Elements Serving as Longevity Determinants/Optimizers Likely Positively Correlates with Organismal Complexity

Complex organisms tend to have more cell and tissue types along with increased specialization in these structures. Cellular interactions are more numerous and more sophisticated. Signaling pathways and their associated biomolecules, though often highly conserved, may utilize additional component derivatives to increase complexity. Due to this sophistication, organisms that are more complex will have more opportunities for problems to occur. Complex organisms require additional "protective" mechanisms to temper this increase in vulnerability, further contributing to increased organismal complexity.

To minimize the severity of the aging phenotype, mechanisms must exist that alter resource allocation and that make other physiological adjustments as the individual ages. These mechanisms must provide a variable and dynamic response, in accordance with an organism's current state of degradation. In organisms with a greater number of biocomponents and potential interactions, additional corrective factors are required to manage these elements and provide an ideal configuration during all phases of life. This logic suggests that the number of longevity determinants and optimizers positively correlates with organismal complexity. On the other hand, this could help to explain how the longevity of simple organisms can benefit significantly from manipulation of only one or a few longevity determinants (Kenyon et al. 1993; Kenyon 2010) while similar manipulations in organisms of greater complexity imparts far more modest longevity benefits (Tatar et al. 2001; Blüher et al. 2003; Selman et al. 2008).

### 10.3 The Rigidity of Species Longevity

Selective pressures have led to highly optimized physiology, based on compromises between factors affecting fitness (e.g. peak biological power density, physical size, longevity, etc.). Consider the potential implications of lowering the mass-specific metabolic rate of a mouse to 1/8^th^ its normal rate (approximately that of a human). The physiology of a murine heart is appropriate for the level of performance of the individual murine cardiomyocytes and the demands of a mouse body. While reducing the metabolic rate by such a large amount would reduce the thermodynamic stresses on cardiomyocytes, it would also mandate a loss in their biological power density. The cardiomyocytes would be less capable of generating the contractile forces necessary to counter the energy dissipation inherent to the murine circulatory system and hence blood circulation would be insufficient for sustaining life. This energy dissipation factor can only be reduced by making fundamental changes to the configuration of the vasculature. Yet, even with a configuration optimized for efficiency over performance, viable metabolic rates will be constrained to those values capable of satisfying all physical requirements. Governing models derived from principles of fluid dynamics have been proposed (West et al. 1997; West and Brown 2005) that provide an example of this type of phenomenon. Although the circulatory system is perhaps the easiest example to conceptualize, there are many other potential negative physiological implications of manipulating singular longevity determinants such as metabolism. A number of other tissues, such as the liver and brain, might be similarly compromised in this type of scenario.

This logic offers an additional, more compelling reason why metabolic manipulations found to increase longevity in simple model organisms translate poorly to organisms that are more complex. While such modifications are well tolerated by organisms such as *Caenorhabditis elegans* (Kenyon et al. 1993; Kenyon 2010), which evidently remain viable at greatly reduced metabolic rates, comparable changes to metabolic rate in mammals are likely to render the organism nonviable. It should be noted that, even in *C. elegans*, reducing metabolic rate via genetic manipulation undoubtedly lowers fitness. This fitness cost explains why *C. elegans* has not evolved to incorporate such changes in their genome: the tradeoff between increased longevity and other impacted factors that affect fitness is not sufficient to select for these changes. For these reasons, I postulate that longevity exhibits a degree of rigidity that increases with organismal complexity and is bound within physiological limits that are explained by fundamental physics.

## 11 Summary and Conclusions

Although evolutionary theory predicts that senescence will arise inevitably due to declining selective pressure with age (Hamilton 1966), fundamental physics stipulates that senescence is inevitable even in the absence of declining selective pressure. Blanket acceptance of declining selective pressure as the *singular* cause of aging is a logical fallacy (converse error)—yet many of the more popular aging theories are grounded in declining selective pressure as the sole root cause. If both declining selective pressure and physics separately stipulate aging, then explaining species longevity requires delineating the respective contributions of each.

### 11.1 Physics Helps to Explain Longevity Trends

It is an interesting observation that organisms with reduced longevity expend more energy on biomolecular repair and replacement. Although this is the precise opposite of what the disposable soma theory of aging would predict, it is in fact quite expected and the logic is straightforward when physics is considered. The model described here predicts that the rate of internal entropy production is greater in organisms with shorter longevity due to increased peak biological power density and that this necessitates upregulated biomolecular renewal to counteract. The result is an increase in the rate of irreversible fidelity loss in susceptible biocomponents. Particularly relevant for many species are the higher thermodynamic stresses placed on DNA molecules (largely through increased replication rates), which lead to an increased rate of loss of DNA informational fidelity and reduced longevity.

This logic offers an explanation for the negative correlation between metabolic rate and longevity, and for the allometry of longevity, which are two of the most long established and irrefutable species longevity trends. Although exceptions to these relationships exist, the model described here also provides rationale for the existence of these exceptions. No theory incorporating declining selective pressure as the primary cause of aging has been able to explain these observations.

### 11.2 Physics Helps to Explain the Hayflick Limit

Leonard Hayflick demonstrated that primary cells undergo a limited number of doublings before senescing (Hayflick and Moorhead 1961; Hayflick 1965), an observation later dubbed the "Hayflick limit". Here it is explained how physics stipulates the loss of DNA informational fidelity and why the degree of loss may be largely a function of replication count. The presence of a replication counter to quantify these losses arises logically from this argument, as does the presence of a replication limit for disposing of cells that have exceeded some threshold of loss. Aging theories based on declining selective pressure are unable to explain why the Hayflick limit exists.

### 11.3 The Degradation State Concept Resolves Previously Unexplained Longevity Paradoxes

Physics clarifies why it is impossible to sustain a biomolecular ensemble in a perfect state of fidelity (because infinite resources are required). The degradation state concept introduced here provides a means to quantify the relative condition of a biomolecular ensemble. A prediction of the described model is that organisms that place extreme priority on maximizing renewal ROI are likely to function at higher degradation states (i.e. with more apparent degradation). It also predicts that organisms with higher peak biological power densities will have lower degradation states at a given life stage, and vice versa. Both of these ideas are counterintuitive, but they help to explain observations previously considered paradoxical and could not be explained by declining selective pressure.

Furthermore, when the inevitability of irreversible fidelity loss in a subset of biocomponents is considered along with the concepts of degradation state and managed deterioration, an explanation emerges as to how and why the overall aging phenotype may develop, including how an organism transitions between new homeostatic states as aging progresses and why many aspects of aging are conserved.

For the above reasons, it is reasonable to propose that stipulations resulting from fundamental physical law are more critical to longevity determination than declining selective pressure. On the other hand, declining selective pressure may help to explain why longevity optimization is less than ideal.

### 11.4 Drawing Incorrect Presumptions from Manipulation of the Aging Phenotype

Typically, the putative aging pathways/genes^9^ have been branded as such because they were found to contain genetic elements which, when altered, modulated longevity in some model organism (often of low complexity, e.g. fruit fly or nematode) or produced a distinct effect on a characteristic(s) typically associated with the aging phenotype. However, fundamental physical law implies that organisms of greater complexity are bound by more stringent physiological restrictions; hence, such manipulations may never deliver substantial longevity benefits to organisms that are more complex. Furthermore, if managed deterioration, as described here, is an actual component of the aging process, then manipulations intended to address specific characteristics of the general aging phenotype will usually carry overall negative repercussions for the individual—regardless of whether or not they "correct" some negative aspect of the aging phenotype—because they disrupt the natural, evolved homeostatic balance that maximizes fecundity/survival at those later life stages.

Observations of singular connections between genetic elements and particular phenotypes demonstrate only that the gene is responsible for modulating those characteristics—it should not imply that the gene is responsible for aging, nor does it necessarily reveal anything about the aging process. This thinking is quite a departure from the current mainstream approach, which focuses on establishing direct relationships between particular genes and their observable effects on the aging phenotype—leading to highly questionable presumptions regarding the culpability of a particular factor as a root-level longevity determinant or “cause” of aging. For these reasons, I believe that the current catalog of putative aging pathways/genes represents, at best, a grossly incomplete set of the factors truly relevant to longevity determination.

### 11.5 Charting a New Course

Aging is the greatest risk factor for severe pathology. Yet, most aging research is focused on age-associated pathologies or metabolic manipulations in lower life forms, rather than the fundamental biology of aging (Hayflick 2000; Hayflick 2007a). Because of this, and despite the fact that a number of pathways influencing longevity have been identified, one can argue that scientists are no closer to a consensus theory of aging today than they were fifty years ago. Even worse, despite the ever-growing litany of serious anomalies challenging common aging theories, the scientific community remains complacent. The multitude of aging theories, the discontinuities between them, and the failure of the scientific community to agree on the root causes of aging, while disappointing, represents a clear opportunity to revisit this problem with a multidisciplinary and somewhat radical approach.

The theoretical framework discussed in this paper utilizes concepts from physics, information theory, as well as evolutionary theory to explain why organisms age and why they live as long as they do. I believe that the theoretical framework arising from this approach has fewer anomalies than existing singularly focused theories. While the concepts put forth here are well supported by the findings of others in diverse fields, it is admittedly not devoid of speculative components. Additional data, such as delineating the species differences in mitobiogenesis rates, would bring better clarity to important questions that remain and would be very helpful in refining the arguments presented here.

The idea that aging can be manipulated is alluring. Nevertheless, difficult problems are only solved by addressing their root causes. In the absence of a central theoretical framework for biological aging, it is hard to predict whether a particular strategy or approach for treating an age-associated pathology, or for increasing healthspan or longevity, has any chance of succeeding in humans. The theoretical framework outlined in this paper provides an explanation as to why the longevity benefits seen in simple organisms through manipulations of putative aging pathways/genes are not realized in organisms that are more complex and why they likely never will be. If accurate, this highlights the naivety of longevity extension efforts to identify and manipulate genes and molecular pathways that could substantially increase human longevity without compromising health or performance, and the futility of attempting to use simple model organisms such as *C. elegans* as the vehicle for such efforts. On the other hand, the capacity for these approaches to extend healthspan in higher organisms (the so-called "intervention potential"), though limited, may be rather straightforward to estimate.

The theoretical framework discussed here identifies fundamental physical law and evolutionary theory as potential root causes of aging. Of course, we cannot manipulate the laws of physics or fabricate exceptions to evolutionary theory, but it may be possible to resolve their most direct downstream negative repercussions on individual viability—irreversible fidelity loss^10^ in susceptible biocomponents—and effectively retard the aging process, reduce susceptibly to age-associated pathologies, or even restore the individual to a more youthful state. For example, while DNA informational fidelity loss may be inevitable, the complete original genetic sequence of an aged individual can be recreated through the collective sequencing of a population of cells from the individual. Hypothetically, it should be possible to artificially produce youthful cells by sorting extant cells according to degree of DNA informational fidelity loss, correcting any genetic errors, resetting stemness or differentiation state if required, followed by propagation and expansion *in vitro*. Reintroducing these cells into an aged individual could produce some of the aforementioned benefits. Although realizing an intervention such as this has a number of technical hurdles, the potential payoffs could be far greater than the indiscriminate approaches currently prioritized which have little to no chance of ever producing tangible results.

## Glossary

**Biological power density:** The volume-specific rate of external biomechanical, biochemical and bioelectric work (power per unit volume) realizable in a specific cell(s), tissue, etc. *Peak* biological power density is the maximum localized value achievable within a particular organism.

**Biomolecular performance:** A measure of the relative ability of a biomolecule to perform its intrinsic biological function.

**Degradation state:** A parameter used to quantify the level of degradation existing within a biomolecular ensemble ("pool" of a given biomolecule).

**Homeostatic shifts:** The concept that an individual organism transitions between different homeostatic setpoints as it ages and that these shifts are typified by increases in biomolecular degradation state, largely resulting from decreases in biomolecular turnover rates.

**Internal entropy production:** The loss of order caused by the nonzero thermodynamic potentials that exist in all nonequilibrium systems. All forms of biomolecular damage result from the production of internal entropy.

**Longevity determinant:** A factor that directly or indirectly influences the basal rate of loss of fidelity in any biocomponent susceptible to irreversible fidelity loss. Longevity determinants define an organism's basal longevity.

**Longevity optimizer:** A genetic element that elicits an effect that generally becomes progressively more dominant with age and that delays the severity, or rate of progression, of the aged phenotype. Longevity optimizers increase longevity beyond the basal longevity specified by longevity determinants.

**Managed deterioration:** The concept that longevity optimizers modulate the deterioration rate of biocomponents so that those of similar importance degrade at comparable rates and/or reach their failure threshold at equivalent ages. Managed deterioration is proposed as the main mechanism by which longevity optimizers realize their longevity extension effects.

**Negative entropy:** Facilitates an increase in the order of a system. In organisms, negative entropy is generated by repair/replacement processes and biomolecular import. Utilized to combat internal entropy production and maintain biomolecular degradation state.

---

**Supp. Fig. 1.**
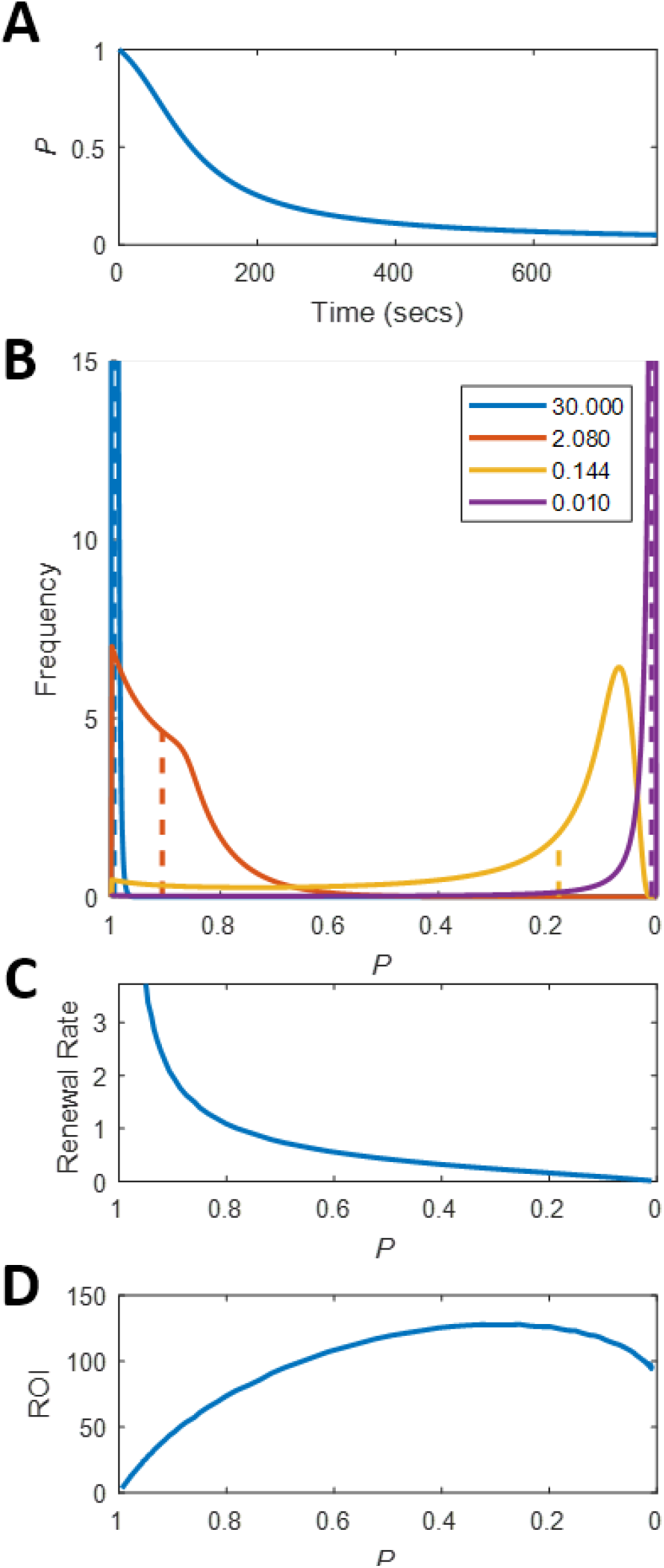
Degradation, renewal and performance characteristics of a hypothetical biomolecule exhibiting excellent stability at high P values. (a) Loss of P with time in a non-renewing population. (b) Simulation of distribution of biomolecules across equivalent P values for different rates of random renewal. Renewal rates are in % of total population replaced per second. Dashed vertical lines indicate P of the ensemble (i.e. average equivalent P). (c) Relationship between P (ensemble) and renewal rate under random renewal conditions. *D* Renewal ROI versus P under random renewal. ROI is in % of *W_m/max_* /renewal rate. Max ROI occurs at P = 0.255.

## Footnotes

1 Not to be confused with free energy minimization during protein folding and the conformational changes of other biomolecules, which occurs very quickly by comparison. In these examples, the free energy minimization involves conformation options for a given atomic structure that can be transitioned to within a short time period. Even in its lowest free-energy conformation, any given biomolecule will still possess considerable excess energy-largely stored in the bonds between its atoms-compared to an equilibrium state where energy dispersion is maximal.

2 Unless otherwise noted, the biomolecules referred to are the proteins, carbohydrates, lipids, nucleic acids, and other macromolecules that define the structure of an organism and facilitate function.

3 As *D* and *P* are ensemble properties, when referring to individual molecules we will prepend the qualifier ‘equivalent’ to acknowledge this and to indicate that the referenced biomolecule's condition is analogous to the typical biomolecule within an ensemble at that given value.

4 Protein synthesis requires approximately 4.5 kJ of energy per gram of protein (Waterlow, 2006, p.170). Producing 245 g/day of protein (human rate) equates to roughly 1103 kJ or 264 Cal per day; 0.09% of this value is 0.23 Cals per day.

5 There are apparently some biomolecules that can accumulate into dysfunctional products when an organism has aged; for example, advanced glycation end products (AGEs), amyloid beta and certain other aggregates (Verzijl et al., 2000). A global decline in biomolecular repair and replacement processes can produce biases leading to significant differences in repair and replacement rates between biomolecules. With infrequent renewal, the proportion of certain types of damaged product can expand, even when youthful renewal rates prevent accumulation. Superficially, this type of damage could be thought of as “accumulated”. An example of this phenomenon was demonstrated by De Baets et al. (2011). There are no published data suggesting that this accumulation occurs under normal circumstances absent significantly decreased renewal such as that which occurs with advanced age. The same proteins found to aggregate with age are produced, but concomitantly cleared, in younger individuals.

6 Although thermodynamics is useful for examining the causes of DNA molecular insults and assessing the magnitude of the damage-inducing potentials, concepts from information theory are more appropriate for analyzing DNA fidelity quantitatively. To avoid confusing the fields, any use of the term entropy in this manuscript refers to thermodynamic entropy. Direct reference to Shannon entropy is avoided.

7 Or from selection occurring on a subcellular level amongst DNA-containing organelles (mitochondria and chloroplasts)

8 The universal value of *f* is contentious. Scientists are largely divided into two camps: one arguing for 2/3 and the other for 3/4 (White and Seymour, 2005).

9 As I subscribe to the belief that aging is a chance-driven catabolic process rather than a genetically engrained behavior (Hayflick, 2007a; 2007b), I view the terms “aging pathways” and “aging genes” as misnomers. I use these terms here only to refer to current literature. “Longevity determinants” and “longevity optimizers” are more appropriate terms for these factors, and this is used when referring to concepts discussed in this paper.

10 It is important to note that “irreversible fidelity loss”, in the context of this discussion, refers to losses that cannot be rectified by any existing, or theoretical, biological process. This does not imply that such losses cannot be resolved through artificial means.

